# Integrated Phosphoproteomics and Transcriptional Classifiers Reveal Hidden RAS Signaling Dynamics in Multiple Myeloma

**DOI:** 10.1101/563312

**Authors:** Yu-Hsiu T. Lin, Gregory P. Way, Benjamin G. Barwick, Margarette C. Mariano, Makeba Marcoulis, Ian D. Ferguson, Christoph Driessen, Lawrence H. Boise, Casey S. Greene, Arun P. Wiita

**Author notes:** **CORRESPONDENCE:** Arun P. Wiita, MD, PhD UCSF Dept. of Laboratory Medicine 185 Berry St, Ste 290 San Francisco, CA 94107.

## Abstract

A major driver of multiple myeloma is thought to be aberrant signaling, yet no kinase inhibitors have proven successful in the clinic. Here, we employ an integrated, systems approach combining phosphoproteomic and transcriptome analysis to dissect cellular signaling in multiple myeloma to inform precision medicine strategies. Collectively, these predictive models identify vulnerable signaling signatures and highlight surprising differences in functional signaling patterns between *NRAS* and *KRAS* mutants invisible to the genomic landscape. Transcriptional analysis suggests that aberrant MAPK pathway activation is only present in a fraction of *RAS*-mutated vs. WT *RAS* patients. These high-MAPK patients, enriched for *NRAS* Q61 mutations, have inferior outcomes whereas *RAS* mutations overall carry no survival impact. We further develop an interactive software tool to relate pharmacologic and genetic kinase dependencies in myeloma. These results may lead to improved stratification of MM patients in clinical trials while also revealing unexplored modes of Ras biology.

## INTRODUCTION

Multiple myeloma (MM) is an incurable malignancy of plasma cells. Considerable effort has gone into deep sequencing of MM primary samples to identify genetic markers for classifying patients into groups for risk assessment and targeted therapy^1–5^. While these studies have offered significant insight into MM biology and prognosis, this knowledge from DNA sequencing largely remains untranslated into evidence-based therapeutic strategies.

We hypothesized that one reason genomic profiles alone have not improved clinical outcome is that they may not be fully predictive of higher-level processes, such as dysregulated cellular signaling, that drive cancer phenotypes. For example, aberrant oncogenic signaling through protein phosphorylation can potentially be inferred, but not directly detected, at the genetic level. Mass spectrometry-based phosphoproteomics has therefore proven a powerful tool to explore cellular-wide signaling alterations at both baseline and under perturbation in cancer^6,7^. For example, studies in acute myeloid leukemia (AML) have shown that phosphorylation signatures can be used to predict increased sensitivity to kinase inhibitors both in cell lines and primary samples^8,9^. Alternatively, one indirect way to assess cellular signaling via a particular pathway is through gene expression profiling. While RNA expression levels of the genes encoding kinases themselves often do not correlate with pathway activation^10^, downstream transcriptional signatures of disparate genes induced by the signaling cascade may reveal specific functional readouts.

In MM, it is thought that aberrant signaling is strongly driven by mutations in the *RAS* family of proto-oncogenes. These mutations are proposed to activate oncogenic signaling primarily via the RAS-REF-MEK-ERK (MAP kinase (MAPK)) pathway and the PI3K-AKT pathway^11,12^. Yet an outstanding mystery in oncology is tumor-specific selection of mutations in different RAS isoforms despite >80% sequence homology and highly similar biological function^12,13^. For example, epithelial-origin tumors predominantly harbor mutations in *KRAS*, whereas *NRAS* mutations are more common in hematologic malignancies, and *HRAS* mutations are rare overall^12^. Furthermore, it is not known why mutations in specific RAS codons are predominantly found in some cancer types (for example, *KRAS* G12D in pancreatic cancer, *KRAS* G12V in lung cancer, or *NRAS* Q61 in melanoma)^14,15^.

MM is a unique case study of Ras biology as approximately 40% of MM patient tumors have predicted activating mutations in *KRAS* or *NRAS*^4,5^, with an almost equal distribution between the two, but essentially none in *HRAS*. Notably, these *KRAS* and *NRAS* mutations are very rare in the precursor lesion monoclonal gammopathy of uncertain significance (MGUS), suggesting that *RAS* mutations are important for transformation to MM^16^. The majority of MM research has treated mutations in these *RAS* isoforms as largely indistinguishable in terms of biological effects, though some studies have begun to elucidate distinctions. For example, clinical outcomes with earlier MM therapies showed patients with *KRAS* mutations had worse overall survival than those with *NRAS* mutations^17,18^. However, later clinical observations suggested that patients with *NRAS* mutations responded more poorly to bortezomib-based therapies than *KRAS*^19^. *In vitro* overexpression studies in myeloma cell lines indicated that *KRAS* may lead to more rapid proliferation in the absence of interleukin-6 (IL-6) stimulation than *NRAS*^20,21^. More recent sequencing studies have suggested that *NRAS* mutations tend to cluster with specific genomic aberrations in MM^5^. Finally, immunohistochemistry studies on MM bone marrow have suggested possible differences in ERK phosphorylation depending on *RAS* isoform and specific mutation^22^. Therefore, this evidence suggests that *KRAS* and *NRAS* in MM are not exactly equivalent, but much about the biology of these differences remains unclear.

Here, we apply an integrated approach using both unbiased phosphoproteomics and machine learning-based classifiers of transcriptional response to dissect signaling in MM. Our results reveal differential kinase activity across MM cell lines with potential implications for selective kinase vulnerability. We next uncover underlying transcriptional output differences for patients with *KRAS* and *NRAS* mutations. Surprisingly, we find that only a fraction of *RAS*-mutated patients are predicted to have highly activated MAPK signaling vs. WT *RAS* patients. However, this group, which is particularly enriched in patients with *NRAS* Q61 mutations, carries the poorest prognosis under modern treatment strategies. Our results identify *RAS*-mutated MM patients who may benefit from precision medicine strategies and reveal modes of *RAS* isoform-driven biology with implications across *RAS*-mutated cancers.

## RESULTS

### Kinase Activity from Phosphoproteomics is Modestly Predictive of Kinase Inhibitor Sensitivity in Multiple Myeloma Cell Lines

Given promising prior results in AML^8^, we first aimed to predict differential kinase activity across MM cell lines and investigate whether increased kinase activity led to increased vulnerability to selective inhibitors. We used immobilized metal affinity chromatography (IMAC) to enrich phosphopeptides across 7 MM cell lines (AMO-1, MM.1S, L363, INA-6, RPMI-8226, KMS-34, KMS-11) as well as one *in vitro*-evolved bortezomib-resistant cell line (AMO1br)^23^ (**Fig. 1A**). Phosphoproteomic profiling was performed in biological triplicate per cell line and in total 19,155 phosphosites from 4,941 proteins were quantified across all cell lines after imputation using MaxQuant^24^ and R^25^ software (**Table S1**). As expected with IMAC enrichment, >99% of all identified phosphorylation events were on Ser or Thr sites, with only a very small minority of Tyr sites. The Pearson *R* of phosphosite intensity correlations between replicates of each cell line ranged from 0.86-0.95, underscoring reproducibility of the phosphoproteomic analysis (**Fig. S1**).

**Figure 1.**
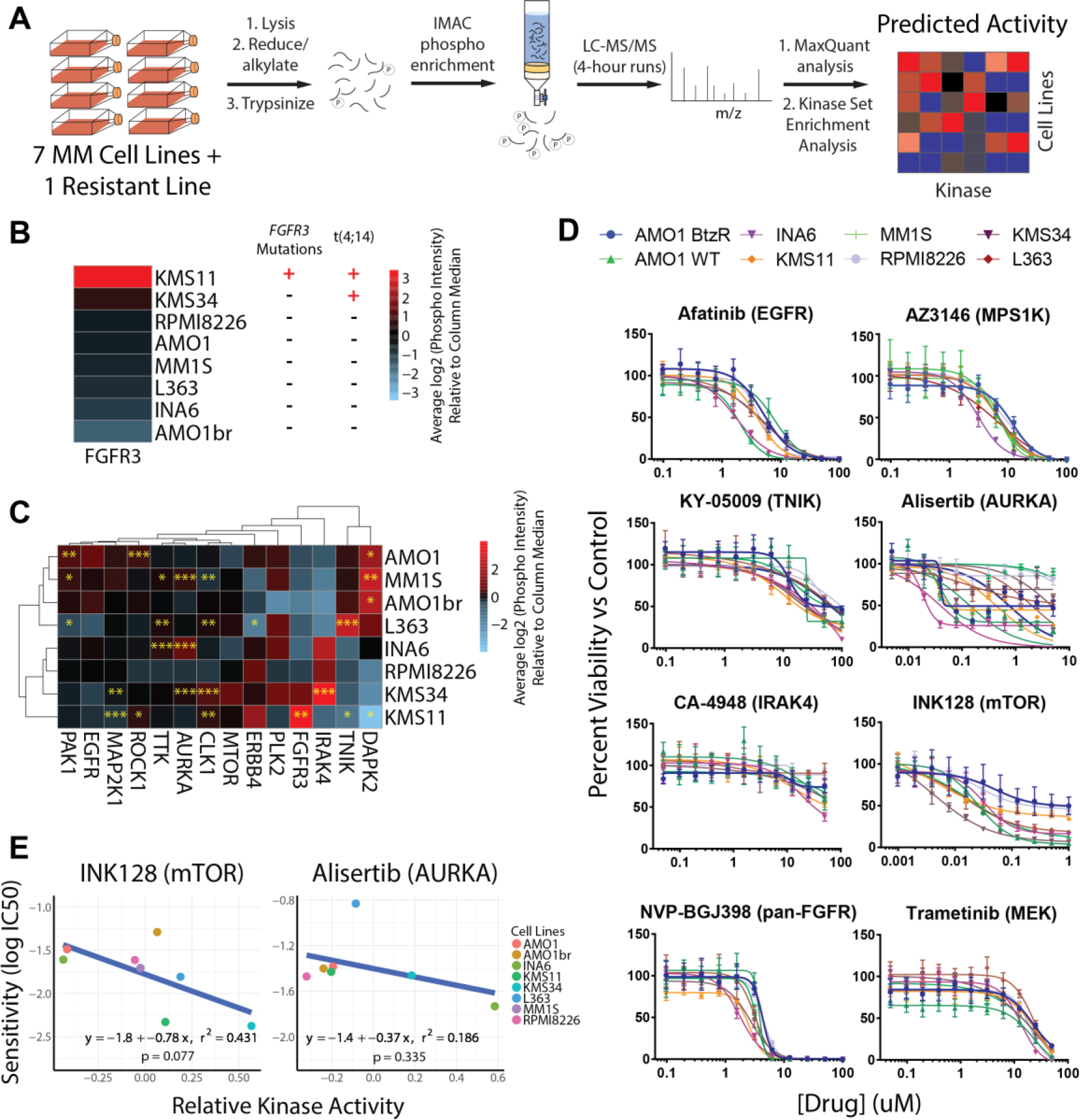
Predicting kinase activity and inhibitor sensitivity in MM by unbiased phosphoproteomics. **A.** Schematic of the pipeline for kinase activity prediction from phosphoproteomic data. All phosphoproteomics were performed in biological triplicate and combined by averaging the log_2_-transformed intensities of phospho-sites associated with each kinase to generate one activity score. **B.** Association of predicted FGFR3 activity from KSEA with known genetic aberrations. **C.** Heatmap of the predicted activities of 14 kinases that exhibited differential activity signatures across myeloma cell lines. The significance of the score from the median activity across cell lines was calculated by z-statistics (see methods). * *p* ≤0.05, ** *p* ≤0.01, *** *p* ≤0.001. **D.** Viability curves showing the drug response of eight myeloma cell lines to eight kinase inhibitors (*n* = 4, mean +/− S.D.), with only INK128 and Alisertib exhibiting strongly differential effects. **E.** Correlation between inhibitor sensitivity and KSEA-predicted kinase activity for mTOR and aurora kinase A across myeloma cell lines shows modest predictive power. *p*-values were calculated based on the null hypothesis that no relationship exists between the activity of a kinase and its sensitivity to an inhibitor.

We first analyzed this data using Kinase Set Enrichment Analysis (KSEA)^26^ to identify kinases with predicted significant differential activity between at least one cell line vs. all others. As an initial validation of KSEA predictions, we found that the KMS-11 cell line, which harbors both a t(4;14) translocation as well as an activating mutation in *FGFR3*, showed the highest predicted activity of FGFR3 kinase (**Fig. 1B**). KMS-34, which has a t(4;14) translocation increasing *FGFR3* expression but no *FGFR3* mutation, showed the second-highest predicted FGFR3 activity. Overall, with the known limitation that KSEA relies heavily on computationally predicted kinase-substrate relationships derived from NetworKIN^27^, we ultimately identified 14 kinases which appeared to have differential activity (**Fig. 1C**) across MM cell lines.

We next evaluated whether these differential kinase activities predicted relative cell viability to selective inhibitors. We were able to obtain the described selective inhibitors against 13 of these 14 kinases. Using an initial pre-screening versus a limited panel of three MM cell lines (AMO-1, MM.1S, RPMI-8226), we found that only eight of these inhibitors demonstrated any anti-myeloma effect at concentrations up to 20 μM (**Fig. S2A**). We tested these eight inhibitors versus the full panel of eight MM cell lines used for phosphoproteomics (**Fig. 1D**). Evaluating sensitivity data, we were surprised to find that only two of the eight inhibitors tested in the full panel (and only two of the 13 total inhibitors tested) showed a notable distribution of LC_50_’s across the tested MM cell lines. We more closely examined these two inhibitors, alisertib targeting Aurora kinase (AURKA) and INK128 targeting the mTOR kinase complex 1 and 2 (mTORC1/2), finding a modest correlation between predicted kinase activity and sensitivity to inhibitor (**Fig. E**). Within the smaller range of LC_50_’s for other inhibitors, we did not find correlations between predicted kinase activity and measured inhibitor LC_50_. However, we do note that KMS-11 and KMS-34 are two of the three most sensitive lines to the pan-FGFR inhibitor NVP-BGJ398 (**Fig. S2B**). Notably we saw no correlation between predicted MEK1 (*MAP2K1*) activity and sensitivity to the MEK1/2 inhibitor trametinib (**Fig. S2B**). Overall, these results suggest that kinase activities from KSEA are modestly predictive of sensitivity to targeted kinase inhibitor therapy.

### Phosphoproteomics Reports on Specific Alterations in MAP Kinase Pathway Activity as a Function of RAS Mutation Status

Given the somewhat limited predictive power observed in the KSEA analysis, we next turned to manual curation of downstream effectors of central cancer signaling pathways. As mutations in *RAS* proto-oncogenes are the most commonly seen single nucleotide genomic alteration in MM^5^, we specifically examined known Ras effectors. Phosphorylation of the three RAF isoforms, ARAF, BRAF, and CRAF/RAF1, is reflective of MAPK activation immediately downstream of RAS^28^. We were intrigued to find that cell lines with *RAS* mutations did not show uniform RAF isoform phosphorylation. Instead, the two cell lines (MM.1S and RPMI-8226) with canonical activating mutations in *KRAS* (both G12A) showed the highest levels of BRAF and ARAF phosphorylation (**Fig. 2A**) In contrast, lines with *NRAS* Q61H and G12D mutations (L363 and the IL-6 dependent cell line INA-6, respectively), as well as the less well-characterized A146T mutation (AMO-1) showed RAF phosphorylation indistinguishable from cell lines with WT *RAS* but potentially increased MAPK activity via FGFR3 alteration (KMS-11 and KMS-34). Similar to findings in other studies^29^, by Western blotting we further found that RAS mutation status did not lead to consistent levels of ERK1/2^T202/204^ phosphorylation, a commonly-used readout of MAPK activity (**Fig. 2B**).

**Figure 2.**
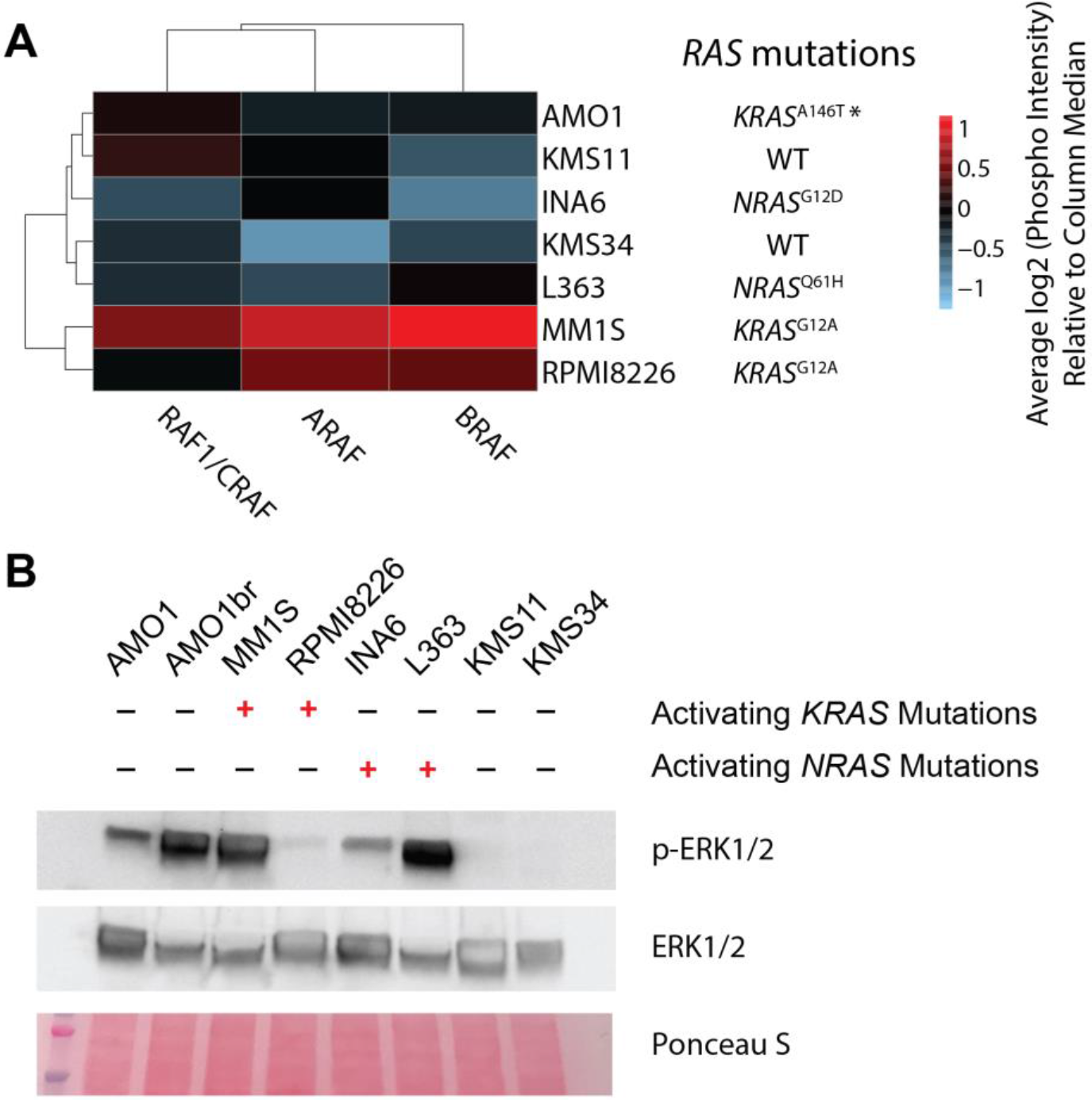
KRAS mutant MM cell lines show greatest activation of downstream RAF effectors based on phosphoproteomics. **A.** Heatmap of phosphoproteome-predicted activity of the immediate downstream substrates of Ras protein (ARAF, BRAF, and RAF1/CRAF) across the eight profiled myeloma cell lines. **B.** Western blot analysis of phospho-ERK1/2^Thr202/204^ and total ERK1/2 levels in MM cell lines, with Ponceau as loading control, demonstrates limited correlation between *RAS* mutant status and the most common biomarker of MAPK activity.

### A Machine-Learning Based Classifier Distinguishes Transcriptional Output of *KRAS* vs. *NRAS* Mutants in MM Patients and Cell Lines

These findings from phosphoproteomics motivated us to further evaluate the biological differences between *KRAS* and *NRAS* mutations in myeloma. Our initial goal was to expand these analyses beyond cell lines to MM patients. As MM patient samples are not typically amenable to phosphoproteomics due to sample input limitations, we turned to more widely available transcriptome data. We recently reported a machine learning-based classifier based on an elastic net penalized logistic regression able to predict *RAS* genotype, or *RAS*-mutant-like phenotype, from tumor RNA-sequencing data^30^. This initial classifier trained and tested solely on solid tumor data from The Cancer Genome Atlas (TCGA) Pan-Cancer Atlas Project, did not distinguish between *RAS* isoforms. Applying this initial classifier to tumor cell RNA-seq data from 812 patients in the Multiple Myeloma Research Foundation (MMRF) CoMMpass study (release IA11a; research.themmrf.org), we observed very limited predictive power for *RAS* genotype (**Fig. S3A**).

Review of gene weights in the initial *RAS* classifier revealed that many highly-weighted genes were transcripts expressed highly in other tissues but minimally in hematopoietic cells (not shown), potentially leading to this lack of applicability. Our initial TCGA classifier was trained using multiple tumor types, but Ras signaling events have previously been detected using machine learning applied to a single tumor type^31^. We therefore used a similar machine learning strategy to build an MM-specific classifier based on CoMMpass patient data. We extended our prior computational approach by developing a three-way classifier, attempting to distinguish transcriptional signatures of patients with WT *RAS*, *KRAS* mutations, and *NRAS* mutations (**Fig. 3A**). For building the classifier, we included patients in the mutation category if any *KRAS*/*NRAS* mutations were reported in CoMMpass data, irrespective of variant allele frequency (VAF). 10 patients with subclonal mutations in both *KRAS* and *NRAS* were excluded.

**Figure 3.**
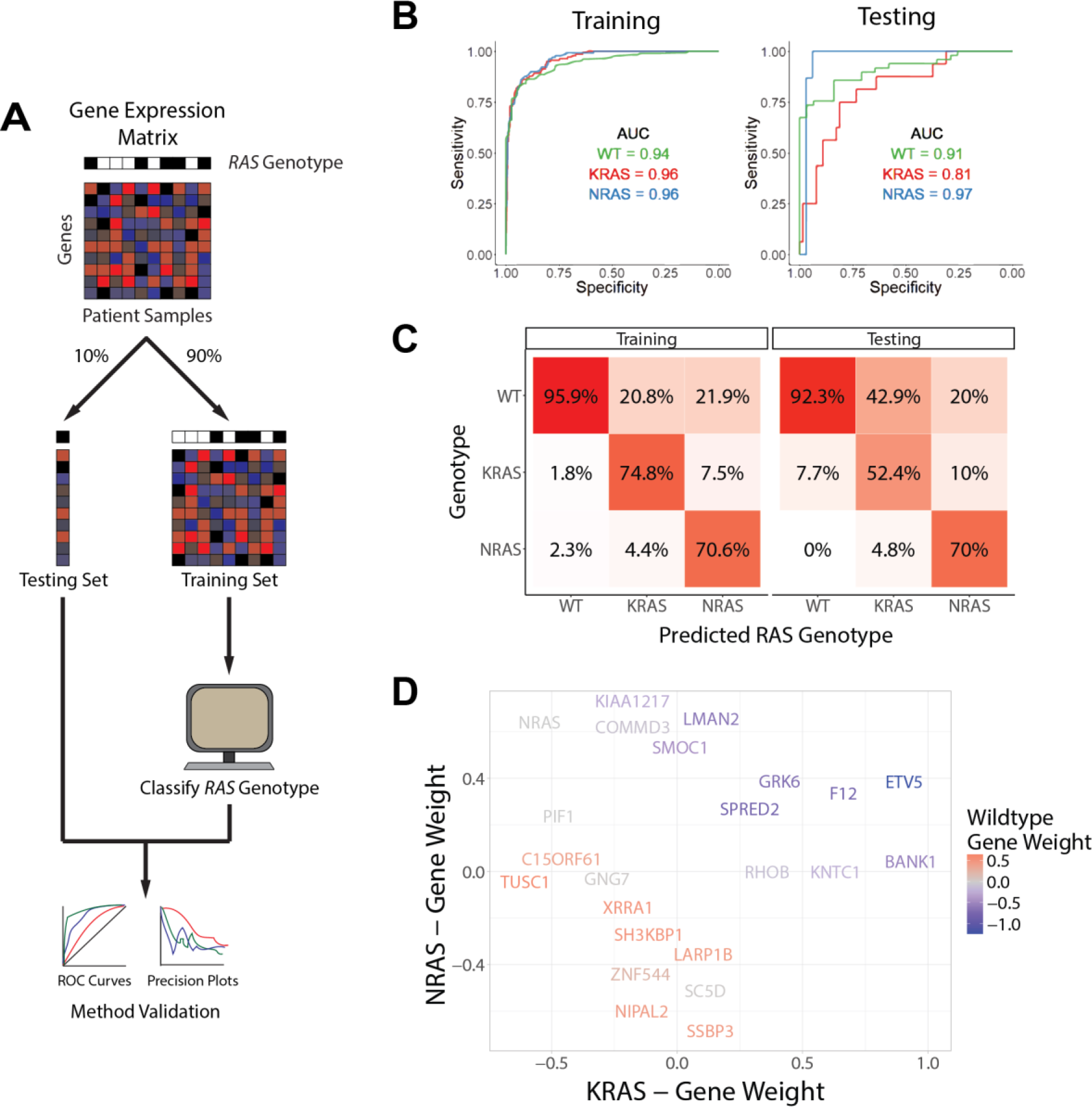
An MM-specific, transcriptome-based *RAS* classifier reveals genes driving the *NRAS* and *KRAS* phenotype. **A.** Workflow for training and testing a gene expression-based machine learning algorithm to predict RAS genotype using an elastic net regression model, adapted from Way et al. 2018. **B.** Receiver operating characteristic curves for evaluating the performance of the predictive model on the training and testing sets. The area under the curve (AUC) is reported for each prediction class. **C.** Confusion matrices showing the fraction of samples in each label-versus-predicted-class combination. **D.** Multi-dimensional plot displaying weighted genes playing the most prominent role in predicting *NRAS*, *KRAS*, or WT *RAS* genotype.

We used 90% of the CoMMpass patient data as a training set (*n* = 706 total; *n* = 439 WT, *n* = 138 *KRAS*, *n* = 129 *NRAS*) and the remaining 10% (*n* = 106 total; *n* = 49 WT, *n* = 16 KRAS, *n* = 15 NRAS) as a holdout test set. The one-vs.-rest, multi-class logistic regression classifier was trained to predict RAS genotype based on transcriptional signatures in the 8,000 genes that exhibited the greatest variance in expression across CoMMpass samples. Our classifier performed robustly in both the training and test sets, with area under the curve (AUC) of the receiver operator characteristic (ROC) curve in the test set of between 0.81-0.97 (**Fig. 3B**). We also applied the classifier to a dataset of 65 MM cell lines characterized by whole exome sequencing and RNAseq (data from www.keatslab.org). The classifier performed similarly well in these data that were previously unseen by the model, which indicates high generalizability (**Fig. S3B**).

We next investigated whether there were differences in predictions between patients carrying *KRAS* and *NRAS* mutations. To address this, we first examined the “confusion matrix”, assessing which genotype was predicted given each of the observed genotypes. Notably, we found that in both the training and test sets, incorrectly-predicted *KRAS*- and *NRAS*-mutant genotype samples were more likely to be predicted as being WT *RAS* rather than mutation in the other *RAS* isoform (**Fig. 3C**), underscoring divergence in transcriptional output between these mutations. We also observed a dependence on clonality: tumors with higher VAF had less accurate classification between *KRAS-* and *NRAS*-mutated tumors while lower VAF led to less accurate classification between WT and *RAS* mutant samples (**Fig. S3C**).

We further examined the highest weighted genes for the *KRAS* mutant, *NRAS* mutant, or WT *RAS* classifiers (**Fig. 3D**). We found a limited set of genes whose expression levels increased the probability of classification as either *RAS* mutant and decreased probability of WT *RAS*: *SPRED2*, *GRK6*, *F12*, and *ETV5*. Of these genes, *SPRED2* and *ETV5* are well-defined as MAPK-responsive genes^32^. Notably, a prior study also found *ETV5* to be one of only three genes consistently overexpressed in *RAS*-mutant MM models^33^. The biological role of these other genes in relation to Ras activation remains less clear. Overall, the genes which specifically defined increased or decreased probability of *KRAS* or *NRAS* mutant classification were largely independent of each other (genes along X- and Y-axes in **Fig. 3D**). Surprisingly, in our machine learning approach the only gene whose expression strongly predicted for *NRAS* over *KRAS* mutant status was *NRAS* itself.

### RAS Mutants Drive Differential Expression of RAS genes and Oncogenic Addiction in Myeloma

This finding of differential expression of *NRAS* transcript driving *NRAS* genotype prediction motivated us to further examine *RAS* gene expression as a function of genotype in MM patients in CoMMpass. Consistent with our machine learning classifier, we indeed found that patients with a detected *NRAS* mutation had significantly increased expression of *NRAS* compared to *KRAS*-mutated samples (**Fig. 4A**). We also saw a less pronounced but still significant reciprocal relationship, where *KRAS* mutant samples had increased *KRAS* expression vs. *NRAS* samples. Indeed, this finding in MM patients is reminiscent of that found in a prior analysis of cell lines in the Cancer Cell Line Encyclopedia, where mutations in one *RAS* isoform were associated with increased expression of that isoform and depressed expression of the other^34^. We further confirmed a similar relationship between *NRAS* mutation and gene expression across MM cell lines, though we did not observe this finding for *KRAS* (**Fig. S4**).

**Figure 4.**
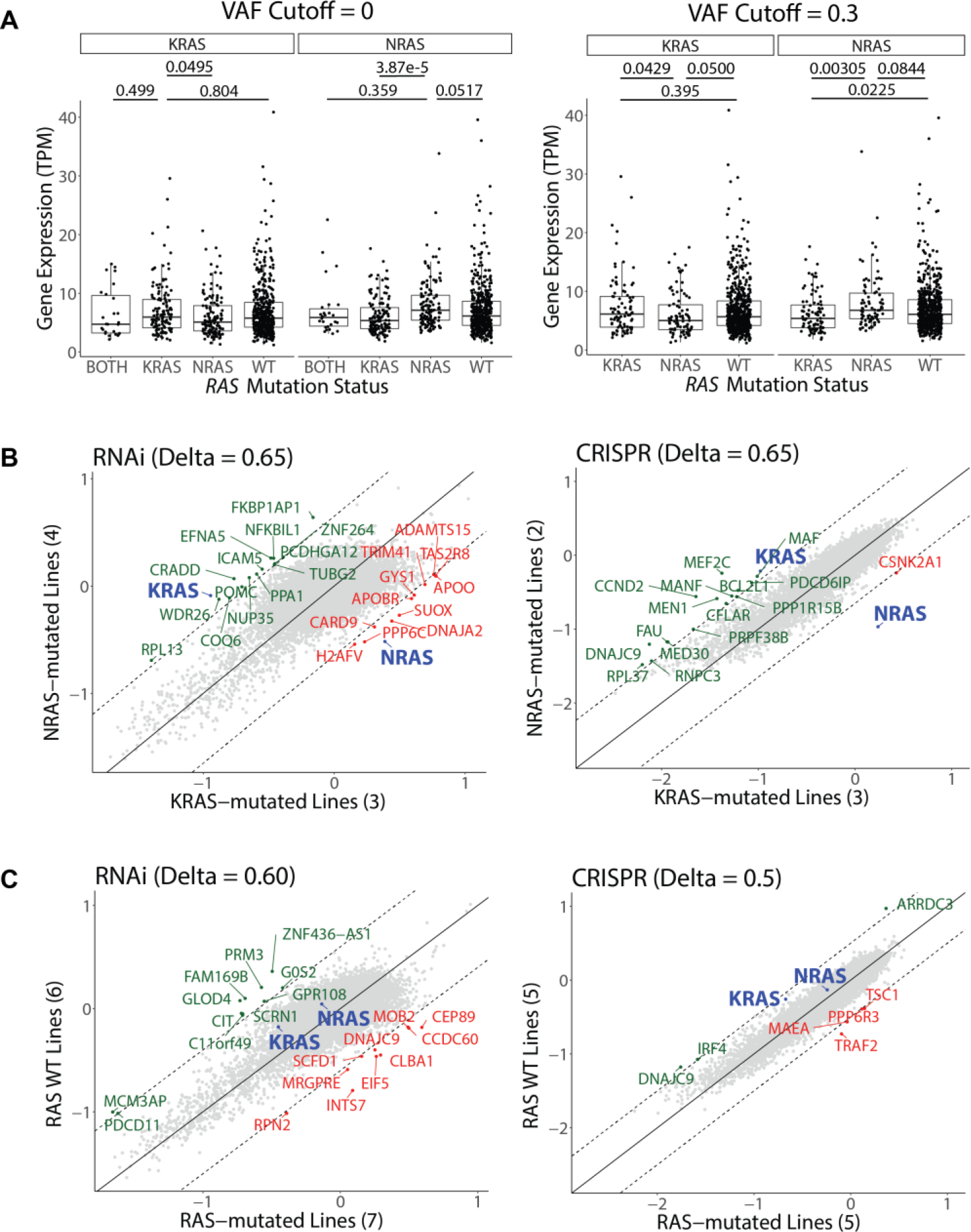
MM-mutant *KRAS* and *NRAS* are associated with differential *RAS* expression and are “addicted” to the mutated *RAS* isoform. **A.** Boxplots of *KRAS* and *NRAS* expression in tumor samples from newly diagnosed MM patients in CoMMpass. The distributions are stratified by *RAS* mutation status and variant allele frequency (VAF) to evaluate their effects on gene expression. *p*-values from Welch’s *t*-tests are reported for relevant comparisons. **B.** Scatterplot of the gene dependency scores from DepMap between *NRAS*-mutated and *KRAS*-mutated MM cell lines from RNA interference (18Q2 release) or CRISPR deletion (Avana 18Q2) functional screens. Comparing across both datasets, only the related *RAS* gene shows consistent essentiality but no other genes. Dashed lines represent cutoffs (Delta) for differential gene dependency between the *RAS*-mutated lines. **C.** Similar to **B**, comparing WT *RAS* and *RAS*-mutated myeloma cell lines.

We next took advantage of data on “essential genes” required for cell viability in the Cancer Dependency Map (depmap.org)^35^. Gene essentiality was determined via genome-wide shRNA knockdown and CRISPR deletion screening across >400 cancer cell lines. We focused on MM cell lines included in this analysis with either an *NRAS* or *KRAS* mutation. Comparing essential genes, we confirmed findings from earlier single-gene knockdowns^36^ that *NRAS*-mutated cells are highly dependent on *NRAS*, and *KRAS*-mutated cells are dependent on *KRAS* (**Fig. 4B**). Surprisingly, though, this genome-wide analysis did not identify any other genes in both the shRNA and CRISPR screen data which led to specific dependency in *NRAS* or *KRAS* mutants, nor between *RAS*-mutated cell lines and WT RAS lines (**Fig. 4C**). These findings underscore the profound “oncogene addiction” of MM plasma cells to mutated *RAS*.

### KRAS Mutations at any Codon and NRAS Q61 Mutations Lead to Poorer Prognosis

We next examined the relationship between *RAS* mutation status and clinical outcomes in CoMMpass. If we included all patients with a detected *RAS* mutation, either at the level of any detected RAS (VAF >0) or those with a dominant subclone (VAF >0.3), we actually found no differential effect on either progression-free survival (PFS) or overall survival (OS) (**Fig. 5A**). This finding suggests that *RAS* mutation status alone does not offer clear prognostic information in MM, consistent with prior studies^37^.

**Figure 5.**
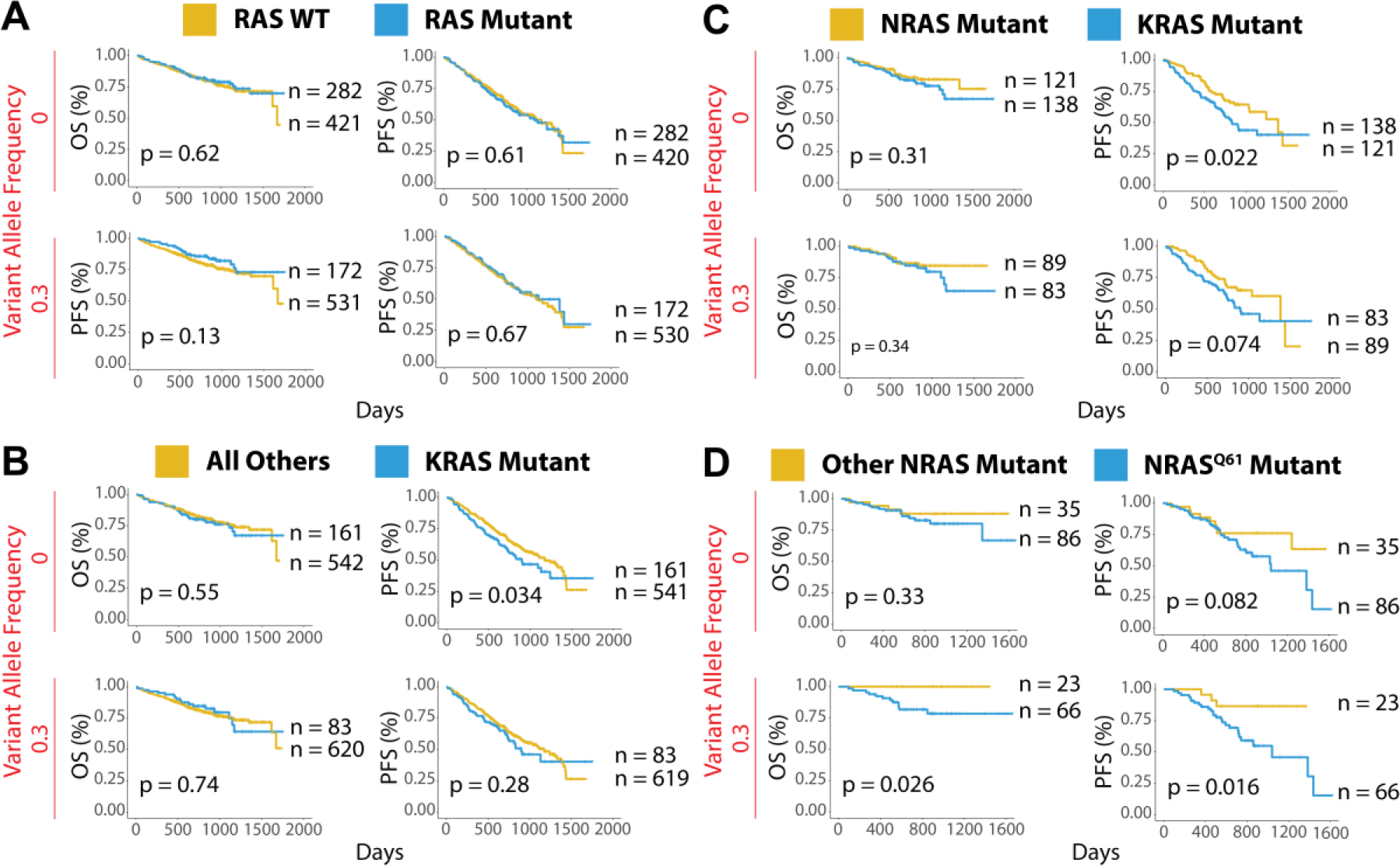
*KRAS* mutations and *NRAS* Q61 predict worse outcomes in MM. **A.** Survival curves comparing the clinical outcome of newly-diagnosed MM patients with and without activating *RAS* mutations (CoMMpass release IA11). **B.** *KRAS*-mutated MM patients vs. all other patients. **C.** Activating *KRAS* versus *NRAS* mutations. **D.** Codon-61 *NRAS* mutations vs. other *NRAS* variants. All analyses were performed at variant allele frequency cutoffs of 0 and 0.3 to assess the effect of tumor heterogeneity on survivorship, with 0.3 as a signifier of a largely dominant RAS-mutated clone for this heterozygous mutation. *p*-values from log-rank test and the sample size for each group are reported. PFS = Progression-free survival; OS = Overall survival.

To further dissect this relationship, next we looked more specifically at effects between *KRAS* and *NRAS* mutations. In this case, we did find that cases with *KRAS* mutation at VAF >0 (i.e. any detectable KRAS mutant reads above background; *n* = 161) did have a significantly decreased PFS vs. all others (*p* = 0.034 by log-rank test), though there was no significant difference in OS vs. all others (*p* = 0.55) (**Fig. 5B**). Surprisingly, if we selected for cases with a dominant *KRAS* mutant subclone (VAF >0.3; *n* = 83), any relationship with survival difference actually disappeared (PFS *p* = 0.28; OS *p* = 0.74).

We similarly evaluated the effects of *NRAS* mutations vs. all others and did not find any outcome effects of *NRAS* mutations overall (**Fig. S5A**). We next compared patients with *KRAS* vs. *NRAS* mutations. Here we again found significantly decreased PFS at VAF >0 (*p* = 0.022) and a similar trend at VAF >0.3 (*p* = 0.074) for *KRAS* vs. *NRAS*-mutated samples, though there was no significant difference in OS (**Fig. 5C**). Overall, these findings appear somewhat consistent with earlier results suggesting that *KRAS* mutations carry worse prognosis than *NRAS*^17,18^, though with modern treatment regimens this poor-prognosis effect of *KRAS* is perhaps not as pronounced.

We next looked at specific effects of mutations in codons 12, 13, and 61 of *KRAS* and *NRAS*; these mutations are most frequently associated with activating mutations in both MM and other cancers^12^. In *KRAS* we did not find significant differential survival effects for mutations in any specific codon (**Fig. S5D-F**). However, in *NRAS* we did note one striking finding: mutations at codon Q61 led to worse PFS and OS vs. other *NRAS* mutations only when present in a dominant subclone (PFS *p* = 0.082 and OS *p* = 0.33 at VAF >0; PFS *p* = 0.016 and OS *p* = 0.026 at VAF >0.3) (**Fig. 5D**). This finding suggests that clonal or near-clonal *NRAS* Q61 mutations are particularly potent in driving disease in the setting of current therapies.

### A Perturbation-based Transcriptional Signature Identifies Highly Variable Activation of the MAPK Pathway across RAS Mutant Samples

Our results thus far strongly support the idea that *NRAS* and *KRAS* mutants in myeloma are not equivalent in driving disease. We next turned our attention to functional readouts of MAPK pathway activation in the context of *RAS* mutation. We took advantage of a recently-described method of transcriptional classification of pathway activity based on perturbation signatures termed PROGENy^38^. This method was validated as providing improved prediction of pathway activation when compared to existing methods of gene expression interpretation^38^. Using the PROGENy prediction for MAPK pathway activity, we were surprised to find a very broad range and prominent overlap of MAPK activation scores across patients with *RAS* mutations versus those with WT *RAS* (**Fig. 6A**). This result stands in contrast to the expected finding of clearly increased MAPK activity in *RAS*-mutated patients vs. WT *RAS*, which, notably, we did find in MM cell lines (**Fig. 6B**). Despite this large degree of overlap in patient samples, at VAF >0 we did still find that the median of the MAPK distribution was statistically significantly increased for both *KRAS* (*p* = 2.16e-6) and *NRAS* mutants (*p* = 0.0123), as well as rare patients with mutations in both genes (*p* = 1.53e-4), vs. WT *RAS*. Overall, our results suggest that many patients with *RAS* mutations do not strongly activate the MAPK pathway over patients with WT *RAS* tumors.

**Figure 6.**
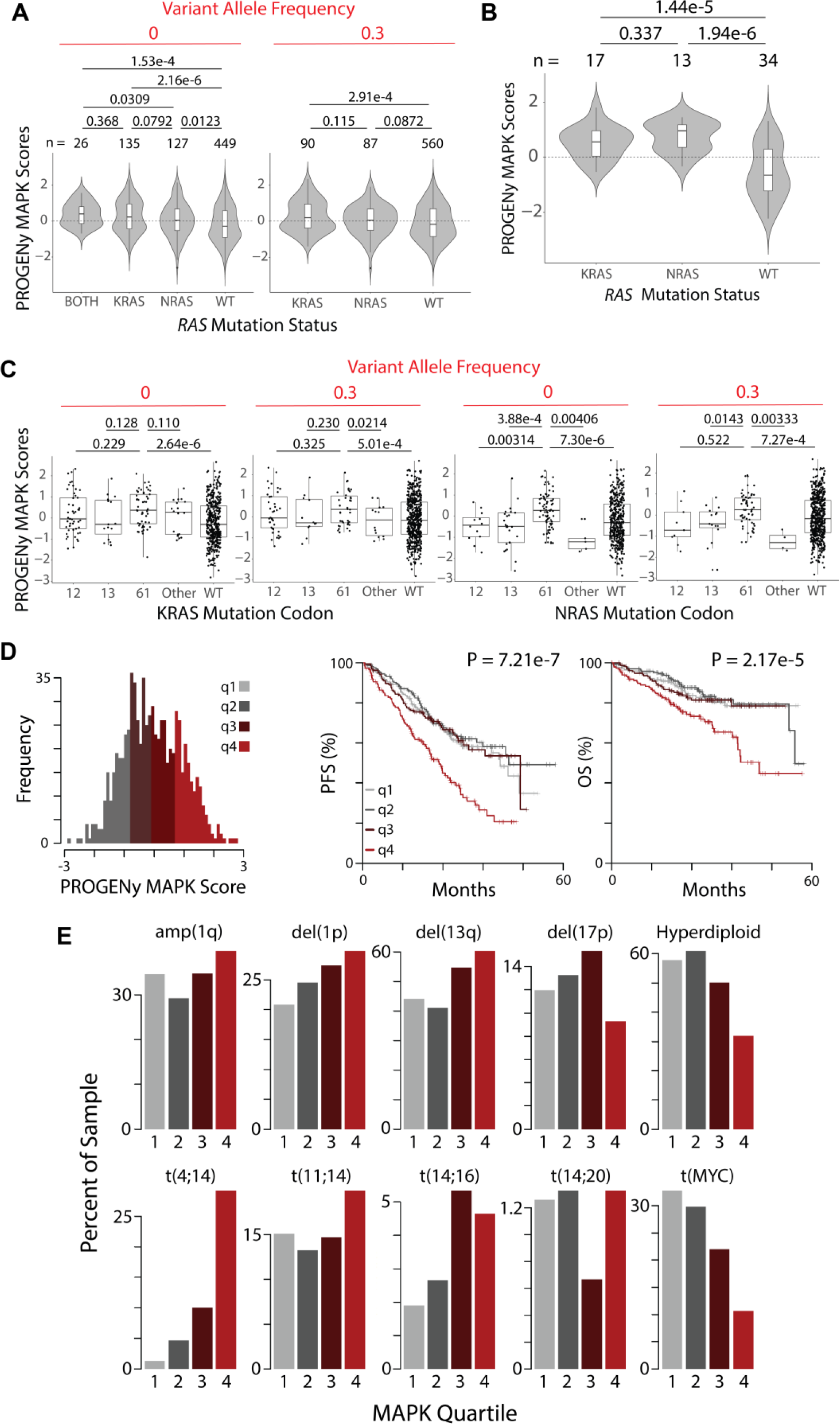
PROGENy reveals RAS mutations do not strongly increase MAPK activity in all RAS-mutant tumors, but patients with increased MAPK activity have decreased survival. **A.** Violin plots showing the distribution of MAPK pathway activation for MM patient samples in CoMMpass based on PROGENy predictions reveals a surprisingly similar range of scores for WT *RAS* and *RAS*-mutated patients, though median of distribution is significantly different. *p*-values for all combinations using Welch’s *t*-tests. **B.** MM cell lines (data from www.keatslab.org) show more pronounced effects of *RAS* mutation driving MAPK activity than patient samples in **A**. **C.** Activating mutations at the Q61 codon show the strongest effect in driving MAPK activity in *NRAS* mutants in CoMMpass samples, whereas *KRAS* mutations do not show similar codon-specific effects. *p*-values by Welch’s t-test. **D.** Histogram of PROGENy-predicted MAPK activation colored by quartiles. Survival analyses demonstrate that high levels of MAPK activity are predictive of poorer outcomes. *p*-values by log-rank test. **E.** Association between PROGENy-predicted MAPK scores and common genomic markers in MM.

Building on our survival analysis in **Fig. 5**, we further examined codon-level effects on PROGENy MAPK score (**Fig. 6C**). For *KRAS*, we found no significant differences between MAPK score for mutations in codons 12, 13, and 61 at either VAF >0 or >0.3. For *NRAS*, in contrast, we found significantly higher MAPK scores for codon 61 mutations vs. 12 and 13. MAPK scores were also markedly higher for codon 61 vs. rare mutations in other non-canonical *NRAS* codons.

### Increased MAPK Activity Predicts Worse Patient Outcomes

These collective observations led us to hypothesize that patients with the highest level of tumor MAPK pathway activity, regardless of *RAS* mutation status, may manifest in more aggressive disease. Consistent with our hypothesis, we indeed found that patients in the highest quartile of MAPK score had significantly decreased PFS (*p* = 7.21e-7) and OS (*p* = 2.17e-5) (**Fig. 6D**).

We further sought to rule out the possibility that increased MAPK score served as a proxy for another known prognostic feature in MM. We did not find any relationship between MAPK score and patient gender, race, age, or β2 microglobulin (**Fig. S6A**). Perhaps expected for a signature of poor prognosis, we did find an association with increased MAPK score and increased M-protein at diagnosis (**Fig. S6A**). We next evaluated the relationship between MAPK score and other MM genomic subtypes (**Fig. 6E**). We do note that we found a significant relationship between MAPK score and t(4;14) translocation frequently associated with FGFR3 overexpression and/or FGFR3 activating mutation^39^. ~ 30% of high MAPK quartile patients carried this translocation whereas it was only present in ~ 1% of patients with MAPK score in the first quartile. This finding is in line with known biology, where FGFR-family tyrosine kinases can activate MAPK upstream of Ras^40^. We also saw more limited associations with other poor prognosis features such as del(1p) and del(13q) (**Fig. 6E and Fig. S6B**). We further evaluated the relationship between MAPK score and common single nucleotide variants and sequence indels present in >2% of CoMMpass patients (**Fig. S6C**). We found a significant association between high MAPK score and both *KRAS* and *BRAF* mutations, but not *NRAS*. Together, these results confirm these MAPK-related genomic lesions as associated with increased MAPK activity, which is consistent with known biology. However, these results also underscore the limitation of genome-only testing: a majority of patients with these well-characterized changes are not in the top quartile of MAPK activity associated with poor prognosis. Overall, our results support the transcriptome-based PROGENy MAPK score as a differential predictor of MM outcomes from other previously known biochemical or genomic markers. This finding also further emphasizes the biological importance of MAPK activity in driving aggressive disease.

### Integrated Analysis of Kinase Activity and Drug Sensitivity Data for Enhanced Precision Medicine in MM

Our results suggest that using targeted therapies specifically for patients with increased MAPK activity, as opposed to *RAS* genotype alone, may be a fruitful strategy in MM precision medicine. To suggest agents which may be most effective in this context, we analyzed data from the Genomics of Drug Sensitivity in Cancer database^41^. Of 265 total compounds tested, we identified three small molecules, the MEK inhibitor refametinib, the MEK inhibitor PD0325901, and the BRAF inhibitor dabrafenib, as showing the greatest correlation (*R*^2^ > 0.65, **Fig. S7**) between LC_50_’s in MM cell lines (*n* = 15) with PROGENy MAPK scores from transcriptome data (**Fig. 7A**). These agents could be potentially considered in clinical applications with targeted use based on tumor transcriptome-based MAPK score.

**Figure 7.**
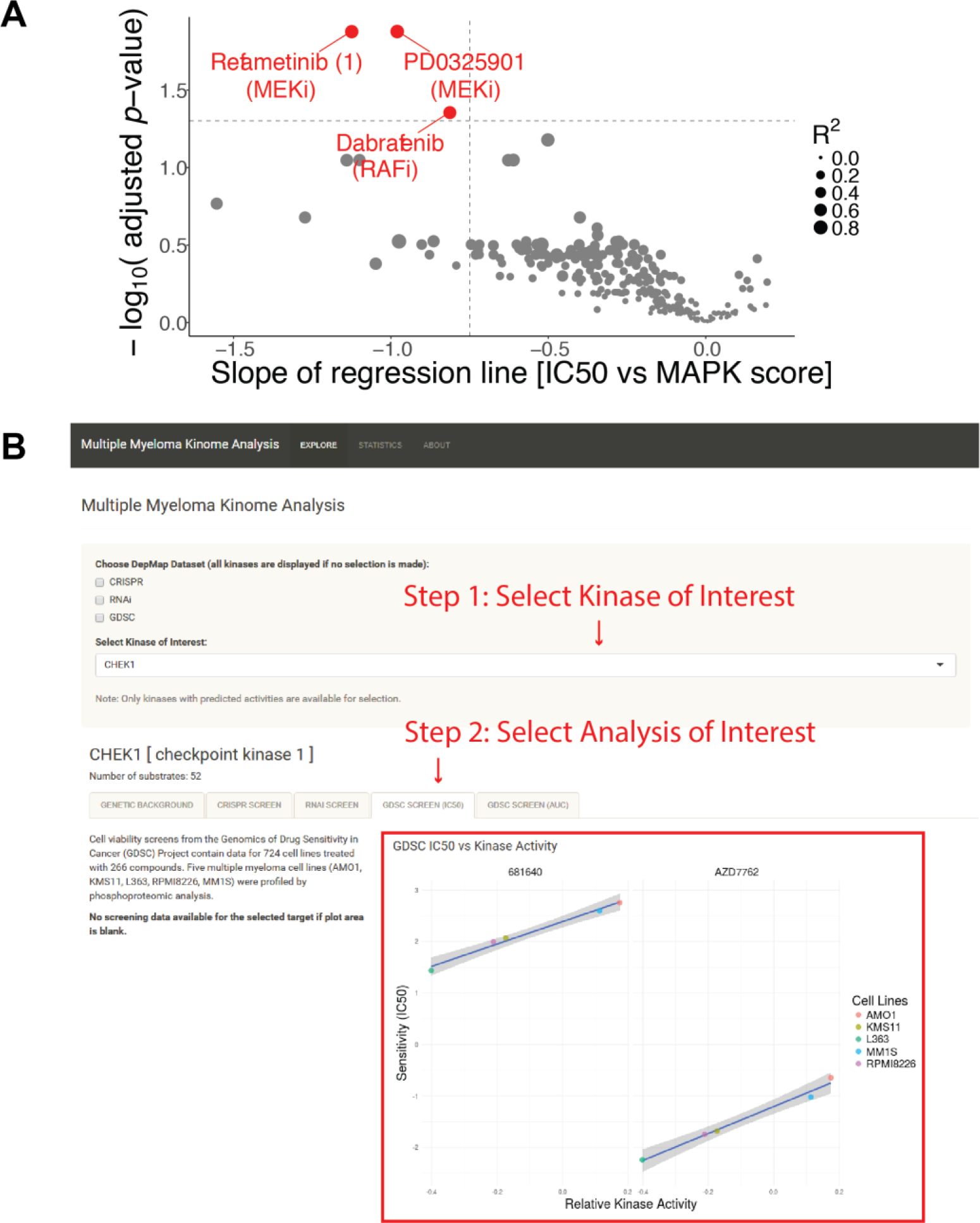
Integration of multiple data types for precision therapy in MM. **A.** Volcano plot showing drug candidates for treating tumors with high predicted MAPK activity. Drug sensitivity data from Genomics of Drug Sensitivity in Cancer database (across 265 compounds tested) and MAPK scores from MM cell line transcriptome data (keatslab.org). *p*-values are calculated based on the null hypothesis that no relationship exists between MAPK activity and inhibitor sensitivity. The three compounds with significant correlations between MAPK activity and drug sensitivity are highlighted in red. **B.** Screenshot of the Multiple Myeloma Kinome Browser (https://tonylin.shinyapps.io/depmap_app/). The example shows the integration of drug sensitivity data with phosphoproteome-based kinase activity predictions on MM cell lines for checkpoint kinase 1.

To assist in further integration of kinase activity into precision medicine, we developed an interactive software tool called the Multiple Myeloma Kinome Browser (MMKB; https://tony-lin.shinyapps.io/depmap_app/) using the *shiny* package (https://shiny.rstudio.com/) in R. This tool combines our phosphoproteome-based kinase activity predictions with multiple data types on MM cell lines from the Cancer Dependency Map (18Q2 release). Our browser integrates functional studies, such as drug sensitivity, CRISPR deletion, and RNA interference screens, and molecular profiles, such as gene expression, copy number variation, and mutation data, for 297 kinases across seven MM cell lines. As one example of its use, we show that a strong, negative association exists between the predicted activity and the sensitivity of checkpoint kinase 1 to two different Chk1 inhibitors (**Fig. 7B**). This integrated, freely-available resource may prove beneficial for future investigation of targeted therapies in MM.

## DISCUSSION

Here we used an integrated approach of unbiased phosphoproteomics and transcriptional classifiers to identify differential regulation of signaling in MM. While differential kinase activity based on phosphoproteomics was only a modest predictor of vulnerability to specific kinase inhibitors, these results suggested downstream signaling differences between *KRAS* and *NRAS* mutated cell lines. This finding led us to further examine *KRAS* and *NRAS* transcriptional signatures in patients in the MMRF CoMMpass study using computational approaches. Our results delineate prominent differences in *KRAS* and *NRAS* signaling outputs, biology, and patient outcomes.

Here, only two of the thirteen kinase inhibitors we tested across a panel of MM cell lines showed any notable differentiation in sensitivity. These two, the mTORC1/2 inhibitor INK128 and the Aurora kinase inhibitor alisertib, showed a modest correlation between predicted kinase activity from KSEA and sensitivity to inhibitor. For INK128 (also known as TAK-228/sapanasertib), we do note that a prior Phase I/II trial in relapsed/refractory MM showed only minimal responses to single agent therapy^42^. Our results suggest that selecting patients based on level of mTOR pathway activation in the tumor may potentially lead to better success rates for other drugs targeting this pathway, such as newly-described bifunctional mTOR inhibitor RapaLink-1^43^. Some of the lack of predictive power of our phosphoproteome analysis is likely biological: simply because a kinase’s activity is increased does not mean that a tumor will be selectively dependent on it for survival. But we are also limited by current bioinformatic tools which are imperfect in predicting kinase activity across thousands of phosphorylated peptides^44^. We were also limited to cell lines, as sample input limitations (requiring >1 mg protein input) currently make probing primary specimens in MM with this technology difficult if not impossible.

Therefore, we focused on our observation that cell lines carrying activating mutations in *KRAS* and *NRAS* demonstrated differential phosphorylation of the immediate downstream MAPK pathway effectors BRAF, ARAF, and CRAF. In studies by others, both in MM cell lines and *ex vivo* primary patient samples, there has been surprisingly little correlation between MAPK activity as measured by ERK phosphorylation and sensitivity to tested MEK/ERK inhibitors^29,45^. We also found no relationship between predicted MEK activity and sensitivity to the MEK inhibitor trametinib from our phosphoproteomic studies (**Fig. 1D**). Immunohistochemical studies of primary patient bone marrow in one study have suggested that the majority of MM patient samples show ERK phosphorylation, regardless of *RAS* mutation status^46^. These findings suggest that ERK phosphorylation alone may actually not be optimal readout to identify MAPK activity driving tumor aggression. Broader-based transcriptional signatures, such as those we derive here, could be more effective in identifying patients who could benefit from therapies targeting this pathway.

Extending from this concept, adapting a machine-learning based classifier of *RAS* mutation status^30^ to the MMRF CoMMpass study revealed new insight into Ras in MM. First, we demonstrate that mutated *NRAS* and *KRAS* are associated with divergent downstream transcriptional signatures in MM. A number of the genes driving the differential classifier have not been canonically associated with Ras, and may reveal new Ras biology in MM. However, one of the strongest predictors of *NRAS* mutant genotype was expression of the *NRAS* gene itself. This relationship between mutation status and gene expression may align with emerging evidence of allelic imbalance across oncogenes, where both mutation status and gene expression converge to drive tumor proliferation^47,48^. These findings further elucidate underlying biological differences between these Ras isoforms in MM that were not previously observed and will underpin further investigation.

Next, we used a recently-described approach called PROGENy that outperforms other existing methods for deriving pathway activation scores from transcriptome data^38^. This analysis is limited in that it relies on computational predictions that cannot be readily validated in a functional manner in this large patient cohort. Nonetheless, we propose that our finding that patients with the highest predicted PROGENy MAPK scores carry the poorest prognosis leads to a pressing question: is there a way to exploit this observation as part of a precision medicine strategy? Our results strongly support the notion that genotype alone is not enough to stratify MM patients. We propose that selecting patients solely based on *RAS* mutation status to receive MEK or ERK inhibitors is unlikely to be effective in MM. Instead, the focus should be on those patients with high MAPK scores, regardless of *RAS* genotype.

While we now have a patient population to focus on, the feasibility of targeting Ras in myeloma remains unclear. We found strongly increased MAPK scores in *RAS*-mutant cell lines vs. WT, but not in patient samples, suggesting the tumor microenvironment *in vivo* may play an important role in modulating MAPK pathway activity. This hypothesis is in line with our prior evidence that IL-6 in the tumor microenvironment strongly modulates MEK signaling and MM cell survival^49^. Furthermore, the high degree of intra-tumoral heterogeneity in MM creates particular hurdles for any therapy that may only eliminate specific disease subclones^4^. However, new approaches to directly target Ras^11^, in addition to existing strategies targeting downstream signaling, may be particularly intriguing in these high-MAPK patients. We further take advantage of PROGENy MAPK scores to leverage broad-scale pharmacologic data. We find that small molecule inhibitors not yet explored in MM show the greatest correlation between MAPK score and sensitivity (**Fig. 7A**). While this analysis is limited as it is only performed in cell lines, it does suggest a functionally-driven, as opposed to genotype-driven, approach to kinase inhibitor selection in MM not previously employed. Notably, our publicly-available tool at https://tony-lin.shinyapps.io/depmap_app/ will enable others to readily evaluate this data to extract other kinase- and signaling-level relationships.

In our analysis, we further identified specific effects of codon-level mutations in MM. In particular, the increased MAPK signaling and poorer outcomes driven by *NRAS* Q61 mutation, but not G12 or G13, are of interest. In terms of biological effects, Q61 mutations in *RAS* (any isoform) have been shown biochemically to activate downstream signaling via complete abolition of GTP hydrolysis and independence of nucleotide exchange factors, as opposed to G12 and G13 mutations which greatly decrease but do not eliminate GTP hydrolysis^12^. This biochemical difference may underpin the greater MAPK scores we find here for Q61, but does not necessarily explain the correlation between higher MAPK activity and poorer outcomes. Murine modeling in melanoma suggests that G12 and G13 *KRAS* mutations are highly effective in driving disease, but for the “weaker” *NRAS* only the “high-output” Q61 mutation is effective enough to strongly promote transformation^50^. Evaluating if similar effects occur in myeloma is an intriguing path forward.

In summary, our results reveal the power of extending genomic studies to the dissection of functional changes within tumor cells using an integrated strategy of phosphoproteomics and computational analysis of transcriptional data. We propose these findings will have broad implications both in MM precision medicine and in the wider study of Ras biology.

## ACKNOWLEDGEMENTS

We thank the Multiple Myeloma Research Foundation for access to the CoMMpass dataset. We thank Drs. Kevin Shannon, Sandy Wong, Nina Shah, Jeffrey Wolf, and Tom Martin for insightful discussions. This work was supported by the Damon Runyon Cancer Research Foundation Dale Frey Breakthrough Award (DFS 14-15), NIH/NCI K08CA184116, NIH/NCI R01CA226851, NIH/NIGMS DP2OD022552, UCSF Stephen and Nancy Grand Multiple Myeloma Translational Initiative (to A.P.W.), MMRF Answer Fund and 5 P30 CA138292 (to L.H.B.), and American Cancer Society Postdoctoral Fellowship PF-17-109-1-TBG (to B.G.B.). G.P.W. was supported in part by a training grant from the NIH (T32 HG000046). This work was funded in part by a grant from the Gordon and Betty Moore Foundation (GBMF 4552) to C.S.G.

## AUTHOR CONTRIBUTIONS

Conceptualization: A.P.W. Methodology: Y.T.L., G.P.W., B.G.B., C.S.G, L.H.B., and A.P.W. Software: Y.T.L., B.G.B., and G.P.W. Formal analysis: Y.T.L., B.G.B., and G.P.W. Investigation: Y.T.L., M.C.M., M.M., and I.D.F. Curation and Resources: Y.T.L. Writing: Y.T.L. and A.P.W. Visualization: Y.T.L., G.P.W., and B.G.B. Funding acquisition: C.S.G., L.H.B., and A.P.W.

## METHODS

### Cell Culture Conditions

All cells were cultured in RPMI-1640 (Gibco, 12-633-012) supplemented with 10% fetal bovine serum (Gemini Bio-products, 100-106), 1% L-glutamine (Corning, 25-005-CI), and 1% penicillin-streptomycin (UCSF, CCFGK003) at 5% CO_2_ and 37°C. INA-6 media was additionally supplemented with 50 ng/mL recombinant human IL-6 (ProSpec, CYT-213). Proteasome inhibitor-resistant AMO-1 was maintained at 90nM bortezomib.

### Phospho-Proteomics Sample Preparation

We profiled eight human myeloma cell lines, RPMI-8226, INA-6, L363, KMS-11, KMS-34, MM.1S, AMO-1, and a bortezomib-resistant AMO-1, by phospho-proteomics. At harvest, 30 million cells were washed twice in room-temperature D-PBS (UCSF, CCFAL001) with 5-minute spins at 300*g* in between each wash. Cell pellets were then flash frozen in liquid nitrogen and stored in −80°C before processing. Replicates were obtained by harvesting cells every two weeks.

We lysed the samples in 6M guanidine hydrochloride (GdnHCl), 0.1M Tris pH 8.5, 5mM TCEP, 10mM 2-chloroacetamide and sonicated the lysate at 1Hz pulses for 45 seconds on ice. Protein concentrations were quantified using the 660nm Protein Assay (Pierce 22660). For each sample, 1 mg of protein was transferred into a clean tube and diluted to 1M GdnHCl by 0.1M Tris pH8.5. Next, we added 20 ug of trypsin (Thermo, PI90057) and incubated the mixture for 20 hours at 37°C.

We halted the trypsin digestion by acidification with trifluoroacetic acid (TFA) to 1% (vol/vol). Any precipitate was removed by centrifugation at 17,200*g* for 5 minutes. The samples were then desalted on a SOLA C18 cartridge (Thermo, 03150391) assisted by vacuum. We washed twice with 0.1% TFA followed by another 2% acetonitrile (ACN)/0.1% formic acid (FA) wash. We eluted the peptides in 80% ACN/0.1% TFA.

Next, we enriched for phospho-peptides using a Fe^3+^-immobilized metal affinity column (IMAC). We prepared the column by stripping the nickel off Ni-charged agarose beads (VWR, 220006-720) using four washes of 100mM EDTA and re-charging them with Fe^3+^ from 150mM FeCl_3_. We then transferred the Fe^3+^-beads to a MicroSpin C18 column (Nest Group, SEM SS18V.25). After IMAC preparation, we transferred our tryptic peptides to the column and performed three washes, under vacuum, with 80% ACN/0.1% TFA to remove non-phosphorylated peptides and two washes with 0.5% FA to equilibrate the C18 resin. To mobilize the phospho-peptides onto C18, we washed twice with 500mM potassium phosphate at pH7. We then performed two final washes with 0.5% FA to eliminate any residual salt. The phospho-peptides were eluted in 50% ACN/0.1% FA, vacuum dried, and stored at −80°C for further analysis.

### Liquid Chromatography-Tandem Mass Spectrometry (LC-MS/MS) Analysis

Enriched phospho-peptides were re-suspended in 2% ACN/0.1% FA. A total of 1.25 ug of peptides from each sample were injected into a Dionex Ultimate 3000 NanoRSLC instrument with a 15-cm Acclaim PEPMAP C18 (Thermo, 164534) reverse phase column. The samples were separated on a 3.5-hour non-linear gradient using a mixture of Buffer A (0.1% FA) and B (80% ACN/0.1% FA). The initial flow rate was 0.5 uL/min at 3% B for 15 minutes followed by a drop in flow rate to 0.2 uL/min and a non-linear increase (curve 7) to 40% B for the next 195 minutes. The flow rate was then increased to 0.5 uL/min while Buffer B was linearly ramped up to 99% for the next six minutes. Finally, we maintained the peak flow rate and Buffer B concentration for another seven minutes before dropping the concentration back to 3%.

Eluted peptides were analyzed with a Thermo Q-Exactive Plus mass spectrometer. The MS survey scan was performed over a mass range of 350-1500 m/z with a resolution of 70,000. The automatic gain control (AGC) was set to 3e6, and the maximum injection time (MIT) was 100 ms. We performed a data-dependent MS2 acquisition at a resolution of 17,500, AGC of 5e4, and MIT of 150 ms. The 15 most intense precursor ions were fragmented in the HCD at a normalized collision energy of 27. Dynamic exclusion was set to 20 seconds to avoid over-sampling of highly abundant species. The raw spectral data files are available at the ProteomXchange PRIDE repository (Accession number PXD011551).

### MS Data Processing

We analyzed the raw spectral data using MaxQuant (version 1.5.1.2) to identify and quantify phospho-sites on serine, threonine, and tyrosine. We relied on default settings except for the inclusion of “Phospho (STY)” as a variable modification. We searched against the Swiss-Prot-annotated human proteome from Uniprot (downloaded August 2016 with 20,163 entries).

We processed the “Phospho (STY) Sites” output file containing the phospho-sites data for downstream analyses in R (version 3.4.0). First, we filtered out proteins annotated as “Reverse” or “Potential contaminant” and retained sites with “localization probability” and “delta score” greater than 0.75 and 8, respectively. To remove poorly quantified phospho-sites, we required each site to be quantified in at least two replicates in one cell line. This reduced the number of identifications from 21,702 to 19,155. We then log_2_-transformed the intensity data and centered the median intensity of each sample at 0 by subtracting the sample median from each value. Missing values were imputed using a hybrid imputation approach, “MLE” setting for values missing at random and “MinProb” for those missing not at random (*imputeLCMD* package (https://cran.r-project.org/web/packages/imputeLCMD/imputeLCMD.pdf)). Finally, we average the phospho-site intensities of the replicates and obtained **Dataset S1**.

### Kinase Set Enrichment Analysis

We implemented the Kinase Set Enrichment Analysis (KSEA) on our MM phospho-proteomics data in R. First, we acquired kinase-substrate associations from a manually curated database called PhosphoSitePlus^51^ (downloaded 2/5/2018 with 10,188 human entries) and from an *in silico* kinase substrate predictive model called NetworKIN^27^ (version 3.0). We downloaded the NetworKIN source code and submitted our phospho-sites data to predict associated kinases using default settings. At a confidence threshold of NetworKIN score >2, we obtained 17,838 associations across 7,256 phospho-sites and 185 kinases. We combined the kinase-substrate associations from PhosphoSitePlus and NetworKIN to obtain 19,328 interactions spanning 7,647 phospho-sites and 297 kinases. Using these annotations, we then inferred kinase activities by averaging the intensity of phospho-sites associated with a given kinase. The significance of the score was determined with a z-statistic, where *mS* is the mean intensity of the sample, *mP* represents the mean of the entire dataset, *m* is the total number of substrates, and *δ* is the standard deviation of the entire dataset.

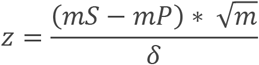

The z-statistic was converted to a one-tailed *p*-value followed by Benjamini-Hochberg correction for multiple hypothesis testing.

### Kinase Inhibitor Screens on Myeloma Cell Lines

To test the hypothesis that the activity of a kinase correlates with its sensitivity, we performed an initial cell viability screen on three myeloma cell lines (AMO-1, MM.1S, RPMI-8226) against 12 small molecule inhibitors of kinases that showed differential predicted activity (**Fig.1C**). We seeded 1e3 myeloma cells per well in 384-well plates (Corning, 07201320) using a Thermo Multidrop Combi and incubated at 37°C, 5% CO_2_ for 24 hours. We then treated cells with PAK1 inhibitor NVS-PAK1-1 (Sigma-Aldrich), TTK inhibitor AZ3146 (Sigma-Aldrich), CLK1 inhibitor TG003 (Selleckchem), TNIK inhibitor KY-05009 (Sigma-Aldrich), ErbB4 inhibitor afatinib (LC Labs), IRAK4 inhibitor CA-4948 (MedChemExpress), PLK2 inhibitor TC-S 7005 (R&D Systems), FGFR3 inhibitor NVP-BGJ394 (LC Labs), MEK1 inhibitor trametinib (ChemieTek), mTOR inhibitor INK128 (LC Labs), ROCK1 inhibitor Y27632 (Selleckchem), AURKA inhibitor alisertib (Selleckchem), or DMSO at 11 concentrations to evaluate dose-response. After 48 hours, cell viability was determined by Cell-Titer Glo reagent (Promega, G7573) using a Glomax Explorer luminescence plate reader (Promega). All drug-dose-cell line combinations were performed in quadruplicates and viabilities were reported as mean (+/− S.D.) ratios normalized to DMSO-treated controls. We used GraphPad Prism 6 for viability curve fitting and to determine the half maximal inhibitory concentrations (IC_50_). This was repeated for a comprehensive screen of our eight MM lines (RPMI-8226, INA-6, L363, KMS-11, KMS-34, MM.1S, AMO-1, a bortezomib-resistant AMO-1) against eight inhibitors (afatinib, AZ3146, KY-05009, alisertib, CA-4948, INK128, NVP-BGJ398, trametinib). The linear fit between IC_50_ values and predicted kinase activity from phospho-proteomics for the eight lines was then computed.

### Western Blot Analysis of ERK

We performed a western blot analysis of total ERK and p-ERK on eight human myeloma cell lines, RPMI-8226, INA-6, L363, KMS-11, KMS-34, MM.1S, AMO-1, and a bortezomib-resistant AMO-1. Immunoblotting was performed using p44/42 MAPK (Cell Signaling Technologies, 137F5) and phosphs-p44/42 MAPK (Cell Signaling Technologies, 9101S) primary antibodies and anti-rabbit IgG secondary antibody (Sigma, NA934VS) according to standard techniques. We stained for total protein as a loading control using Ponceau S (Thermo, BP10310).

### CoMMpass Data Processing for Machine Learning Classifier Input

We downloaded the RNAseq data from CoMMpass (IA11a build) as FPKM estimated by Cufflinks. We also downloaded the somatic mutation file containing canonical variant calls from the CoMMpass website (IA11a build). A total of 812 predicted samples were profiled for somatic variations. Our 812 samples were labeled as wildtype (WT) *RAS* (n = 488), *KRAS* mutants (n = 154), *NRAS* mutants (n = 144), or dual *RAS* mutants (n =26) based on the presence of canonical activating mutations in *RAS* isoforms at codon 12, 13, or 61. To avoid confounding the effects of *KRAS* and *NRAS* mutations on downstream transcriptional programs, dual *RAS* mutants were omitted. We split 90% of the remaining data into a training set (n = 706), which we used to optimize and train the machine learning classifier. We held out 10% of the data (n = 80) as a test set to evaluate performance. The data was split to ensure balanced representation of total *KRAS*, *NRAS*, and WT Ras samples. We z-score transformed the training data by gene to obtain a standardized measure of expression across the 58,095 measured genes and 706 patient samples. To reduce computational burden while maximizing the genes most likely to contribute to classifier performance, we subset the genes in the training set using the top 8,000 most variably expressed genes by median absolute deviation (MAD). These MAD genes were used in all downstream analyses.

### Evaluating the performance of a pan-cancer *RAS* classifier for predicting *RAS* pathway activation in myeloma tumor samples

We applied our elastic net penalized logistic regression classifier for predicting *RAS* pathway activation trained on solid human tumors from The Cancer Genome Atlas (TCGA) PanCanAtlas Project^30^ to the myeloma patient gene expression samples from the CoMMpass study. To apply the model, we first z-scored the full RNAseq CoMMpass data by gene. We then applied the classifier to the CoMMpass data using the following operation:

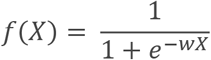

Where *X* represents the CoMMpass scaled RNAseq data and *w* represents the coefficients derived from Way et al. 2018. We determined the status labels as Ras mutant (*KRAS* or *NRAS* mutations) and Ras WT. We evaluated performance of the TCGA classifier applied to the CoMMpass data using a receiver operating characteristic (ROC) curve. Classifier activation scores were tested between groups using a Welch’s t-test.

### Training and evaluating a multiple myeloma-specific machine learning classifier to predict*RAS* genotype

We trained a supervised multi-class machine learning classifier to predict *RAS* genotype using RNAseq data on CoMMpass patients (IA11a build). We used a one-versus-rest (OVR) approach to predict *KRAS* and *NRAS* mutations separately. Specifically, the OVR trained three classifiers: *KRAS* vs. *NRAS* and WT, *NRAS* vs. *KRAS* and WT, and WT vs. *NRAS* and *KRAS*. We trained the models using the sklearn.linear_model.LogisticRegression class (version 0.19.1) in Python 3.5. We searched over a hyperparameter grid of regularization penalties ([0.001, 0.2, 0.4, 0.45, 0.5, 0.55, 0.6, 1, 10]) and loss penalties ([l1, l2]) and selected optimal regularization (C = 0.45) and penalty (L1) terms using a five-fold cross-validation procedure. The final OVR classifiers contained 834 genes with non-zero coefficients.

We evaluated classifiers using ROC and precision recall (PR) curves and calculated the area under the ROC (AUROC) and PR (AUPR) curves for training and holdout test sets. Additionally, we randomly shuffled the gene expression data by gene and evaluated performance of the OVR classifier. This shuffling approach serves as a negative control and will identify potential inflated performance metrics.

We also tested the ability of the multiple myeloma specific classifier to generalize to a cell line dataset. We first processed the expression data (in FPKM) by scaling the genes by z score. Of the 834 non-zero genes, 809 were also quantified in the cell line dataset (97%). We applied the classifier to RNAseq data on 65 MM lines (www.keatslab.org) as described above. We then applied the model and assessed the performance of the MM-specific *RAS* classifier using ROC and PR curves. High performance in the cell line dataset indicates that our OVR classifiers generalize to data previously unseen by our model.

All software to reproduce the analysis and results of our supervised machine learning analysis is provided at https://zenodo.org/record/2566059#.XHTWgeJKiAM.

### Survival Analysis on CoMMpass Patients

All survival curves were based on patient survival data from the CoMMpass study (IA11a build) and were constructed using the *survival* (https://CRAN.R-project.org/package=survival) and *survminer* (https://CRAN.R-project.org/package=survminer) packages in R. We excluded patients who have exited the program prematurely (n = 877 remaining). Survival data were intersected with genetic variant calls on samples from newly diagnosed patients. We label samples that harbor canonical activating mutations in the codons 12, 13, or 61 of any *RAS* isoform as *RAS* mutants. Survival curves were stratified by *RAS* mutation status, and the *p*-values were determined by log-ranked tests.

### MAPK Pathway Activity from Gene Expression Data on Myeloma Cell Lines and Patients

To infer MAPK pathway activity from gene expression data, we applied a perturbation-response machine learning model called PROGENy on RNAseq data from myeloma cell lines (www.keatslab.org) and patient samples (CoMMpass IA11a build). We used the gene counts data as input and performed variance stabilizing normalization using the *DEseq2* library^55^ in R. We then computed PROGENy pathway scores using the *progeny* package (https://github.com/saezlab/progeny). Finally, MAPK scores were z-scored to center the activity values at 0 with a standard deviation of 1 across all samples, separately for cell lines and patients.

The predicted MAPK activity scores can be associated with drug sensitivity data from myeloma cell lines in the GDSC or *RAS* mutation status and survival data from myeloma patients in the CoMMpass study.

### Analysis of CoMMpass MAPK PROGENy Score

CoMMpass clinical and summarized molecular data (interim analysis 11) was provided by the MMRF as part of the participating site agreement. Raw molecular data was downloaded from dbGaP (phs000748.v6.p4), and was approved by the data access use committee. The MAPK PROGENy score quartile was compared to patient PFS and OS using a cox proportional hazards regression using the R / Bioconductor (v3.4.3)^56^ *survival* package (v2.41-3) where p-values were calculated using a Wald’s test. MAPK PROGENy score quartile was compared to patient clinical characteristics (age, gender, race, β2M, and M-protein) using a linear regression for continuous variables and Fisher’s exact test for discrete variables. Common translocations and copy number alterations were determined as previously described^57^. Briefly, translocations were called with DELLY (v0.7.6)^58^ and underwent a number of quality control steps to remove low confidence calls resulting from sequencing reads with high homology or low mappability. Copy number alterations were determined using tCoNut (https://github.com/tgen/tCoNuT) and were defined as having an absolute log_2_ CNA ratio of myeloma DNA to normal DNA of ≥0.2.

### Data Integration with Cancer Dependency Map (Multiple Myeloma Kinome Browser)

We leveraged the high-throughput functional genomic and drug screening data from the Cancer Dependency Map (depmap.org) to construct a web application for exploring kinase dependency in MM. We downloaded the RNA interference^52^, CRISPR deletion (Avana)^53^, and the Genomics of Drug Sensitivity in Cancer (GDSC)^54^ drug screen data from the 18Q2 DepMap release. In addition, we downloaded data on gene copy number, expression, and mutation on DepMap cell lines. The Multiple Myeloma Kinome Browser integrates data on 297 kinases with phospho-proteome-predicted activities across seven human myeloma cell lines, RPMI-8226, INA-6, L363, KMS-11, KMS-34, MM.1S, and AMO-1. We visualized the linear relationships between the sensitivity of a kinase to genetic or pharmacological perturbation and its predicted activity in our myeloma lines. We built the application using the *shiny* library (https://CRAN.R-project.org/package=shiny) in R.

## SUPPLEMENARY TABLES AND DATASETS

**Supplementary Table 1: Summary Statistics on Phospho-proteomic Data (attached as Excel file)**

Eight human myeloma cell lines, including one bortezomib-resistant AMO-1, were profiled in triplicates by mass spectrometry-based phospho-proteomics as described in the methods. The summary statistics, including percent of phospho-sites quantified out of the total identified and percent of phospho-sites imputed for each sample, are reported.

**Supplementary Dataset 1: Quantitative Phospho-Site Data on Myeloma Cell Lines (attached as CSV)**

Mass spectrometry-based phospho-proteomic profiling identifies 19,155 phospho-sites from 4,941 proteins across eight human myeloma cell lines after data processing. The normalized, log_2_-transformed intensities (columns starting with “LOG2”) are provided here. Imputed values are annotated in the categorical columns that start with “IMP”.

**Supplementary Dataset 2: Coefficients of Myeloma-Specific***RAS* **Classifiers (attached as CSV)**

We trained a multiple myeloma-specific machine learning classifier to predict *RAS* genotype using RNAseq data from CoMMpass patients. The classifier weights for the expression of 8,000 genes used in the logistic regression model are presented.

**Supplementary Dataset 3: Top 100 Recurrently Mutated Genes in CoMMpass Patients (attached as CSV)**

A list of the most frequently mutated genes, involved in missense, frameshift, stop-gained, stop-lost, or start-lost variantions, in newly-diagnosed CoMMpass patients is compiled. The differences in PROGENy-predicted MAPK scores between the mutated and non-mutated groups along with the *p*-values (by Welch’s t-test) and false discovery rates (from Benjamini-Hochberg correction) are used to construct **Fig. S6C**.

**Figure S1.**
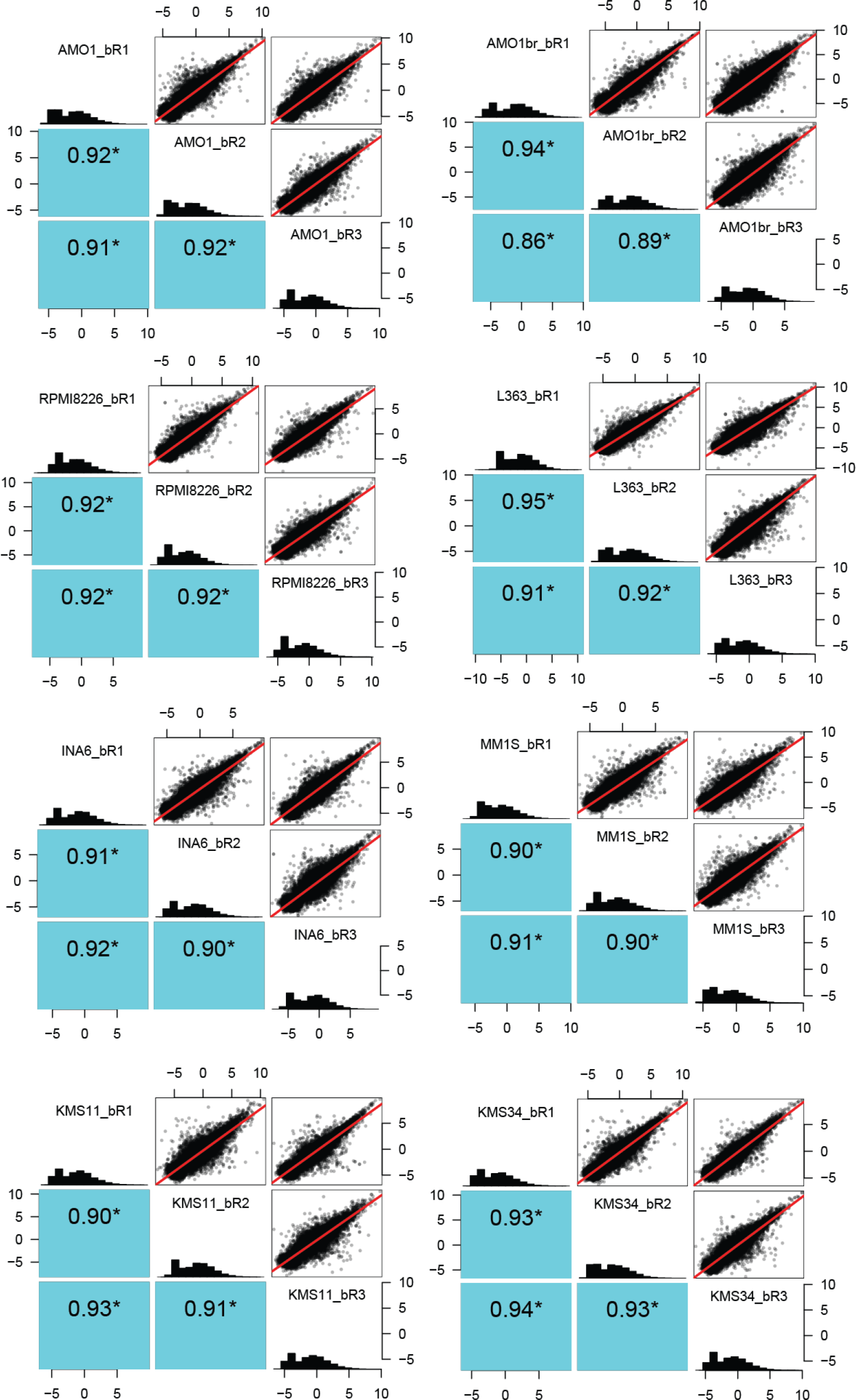
Reproducibility of the phospho-proteomic analysis of MM cell lines. Multi-plots show high reproducibility of phospho-proteomic data from sample replicates. Pairwise scatter plots along with the linear fits (in red) are displayed at the top-right corner. Histograms showing the distribution of phospho-site intensities are found along the diagonal. The Pearson correlation for each pairwise comparison is displayed at the bottom-left corner.

**Figure S2.**
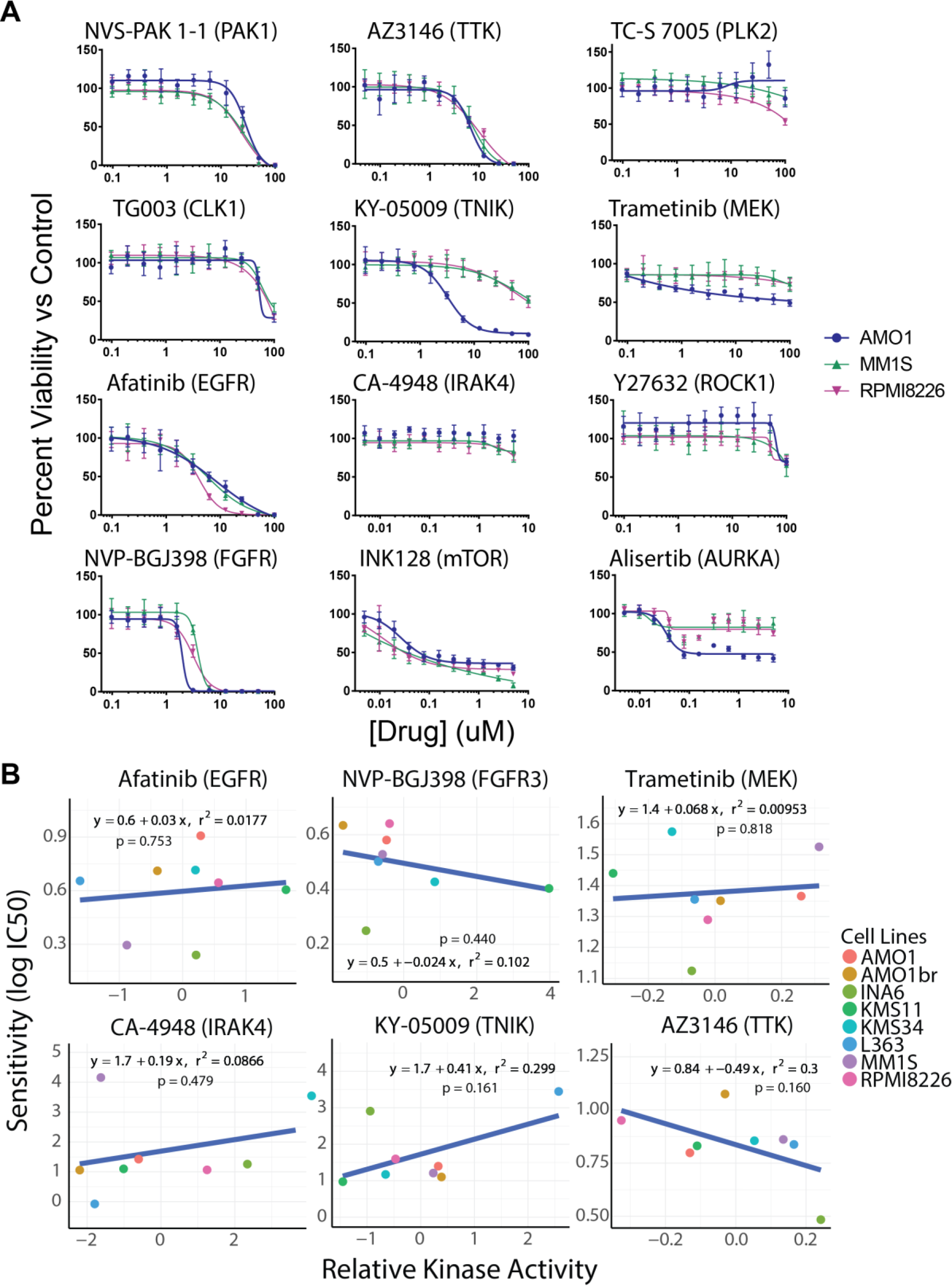
Preliminary drug screening data and associations with predicted kinase activities. **A.** Viability curves showing the drug response of three myeloma cell lines to 12 kinase inhibitors (*n* = 4 +/− S.D.). Eight inhibitors that showed differential responses were screened with the full panel of eight MM cell lines in **Fig 1D**. **B.** Correlation between inhibitor sensitivity and KSEA-predicted kinase activity for six kinases across myeloma cell lines shows low predictive power.

**Figure S3.**
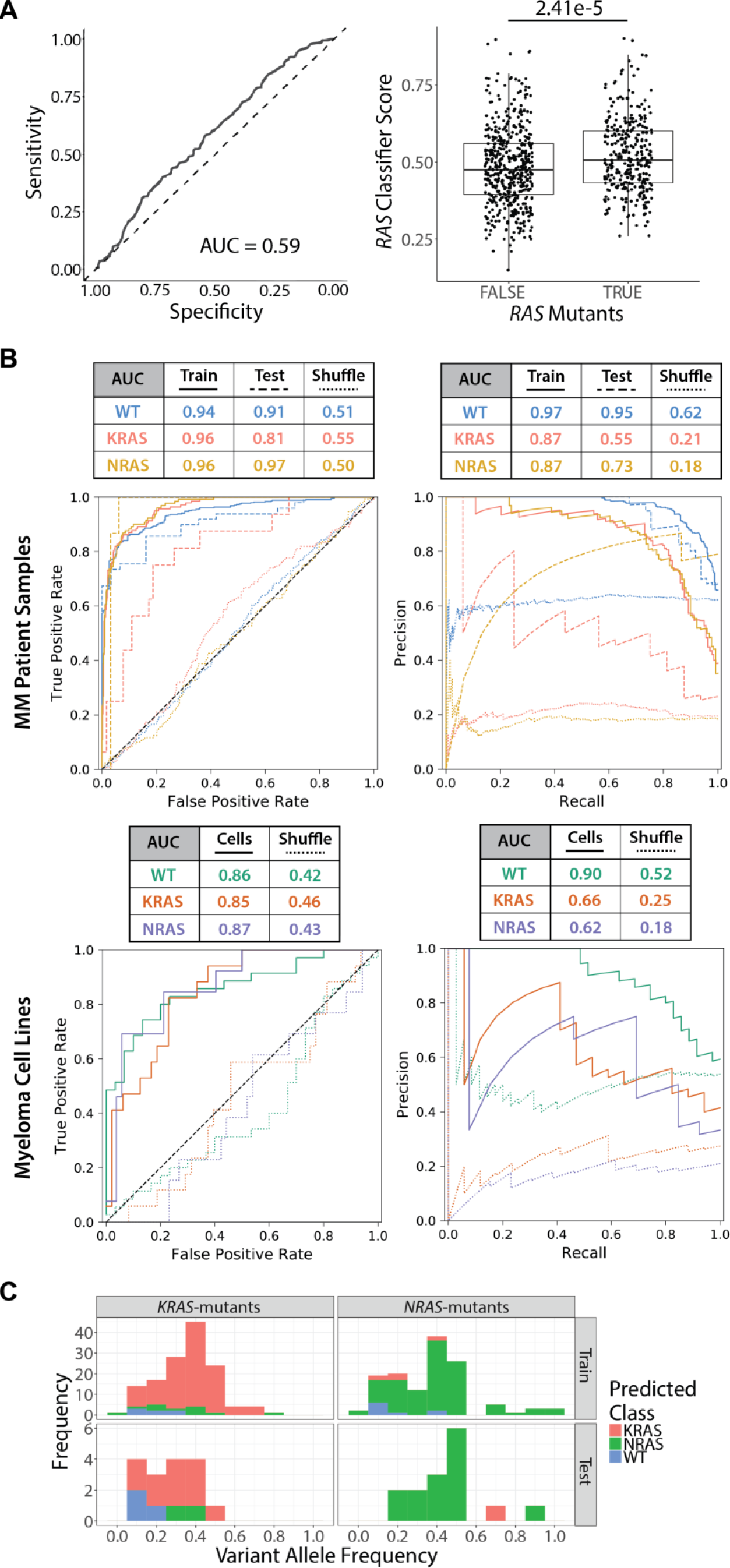
Performance of machine learning classifier for *RAS* genotype prediction. **A.** Receiver operating characteristic (ROC) curve shows relatively poor performance of a pan-cancer Ras pathway activation model (Way et al. 2018) in predicting *RAS* genotype in MM tumors (*left*), though the predicted Ras activation, on average, is higher for *RAS*-mutated samples compared to WT RAS tumors. Statistical significance was determined by Welch’s t-test. **B.** ROC curves and precision-recall plots show much-improved performance of the myeloma-specific transcriptome-based model for predicting *RAS* genotype in MM patients and cell lines under both the training and test setting. **C.** Distributions of the variant allele frequencies (VAF) in *KRAS*-mutated and *NRAS*-mutated samples indicate that *RAS* mutants are misclassified as WT at low VAF but are misclassified as the other *RAS* mutant at high VAF.

**Figure S4.**
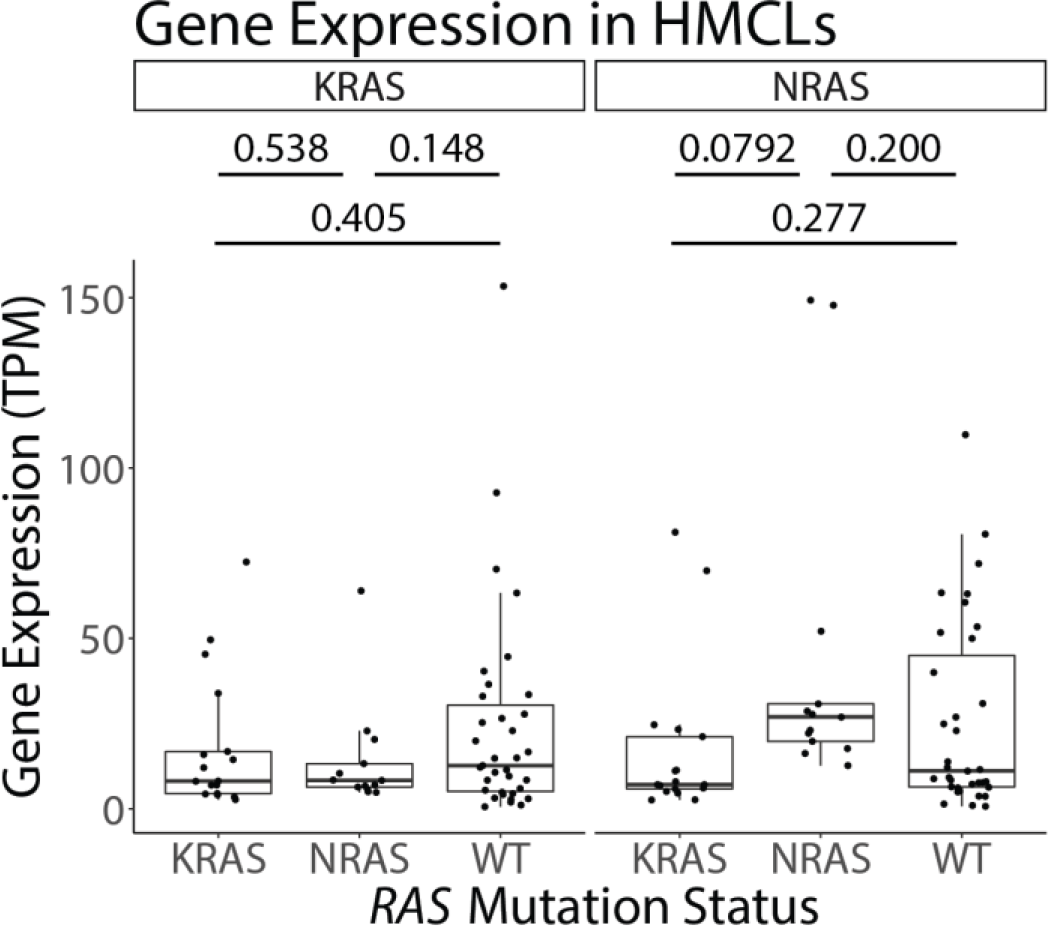
Expression of *KRAS* and *NRAS* in myeloma cell lines. Distributions of the expression of *KRAS* (left) and *NRAS* (right) in MM cell lines show that mutations in *NRAS* is associated with greater *NRAS* expression while the same relationship is absent in *KRAS*-mutated lines. *p*-values are derived from Welch’s t-tests between all sample combinations.

**Figure S5.**
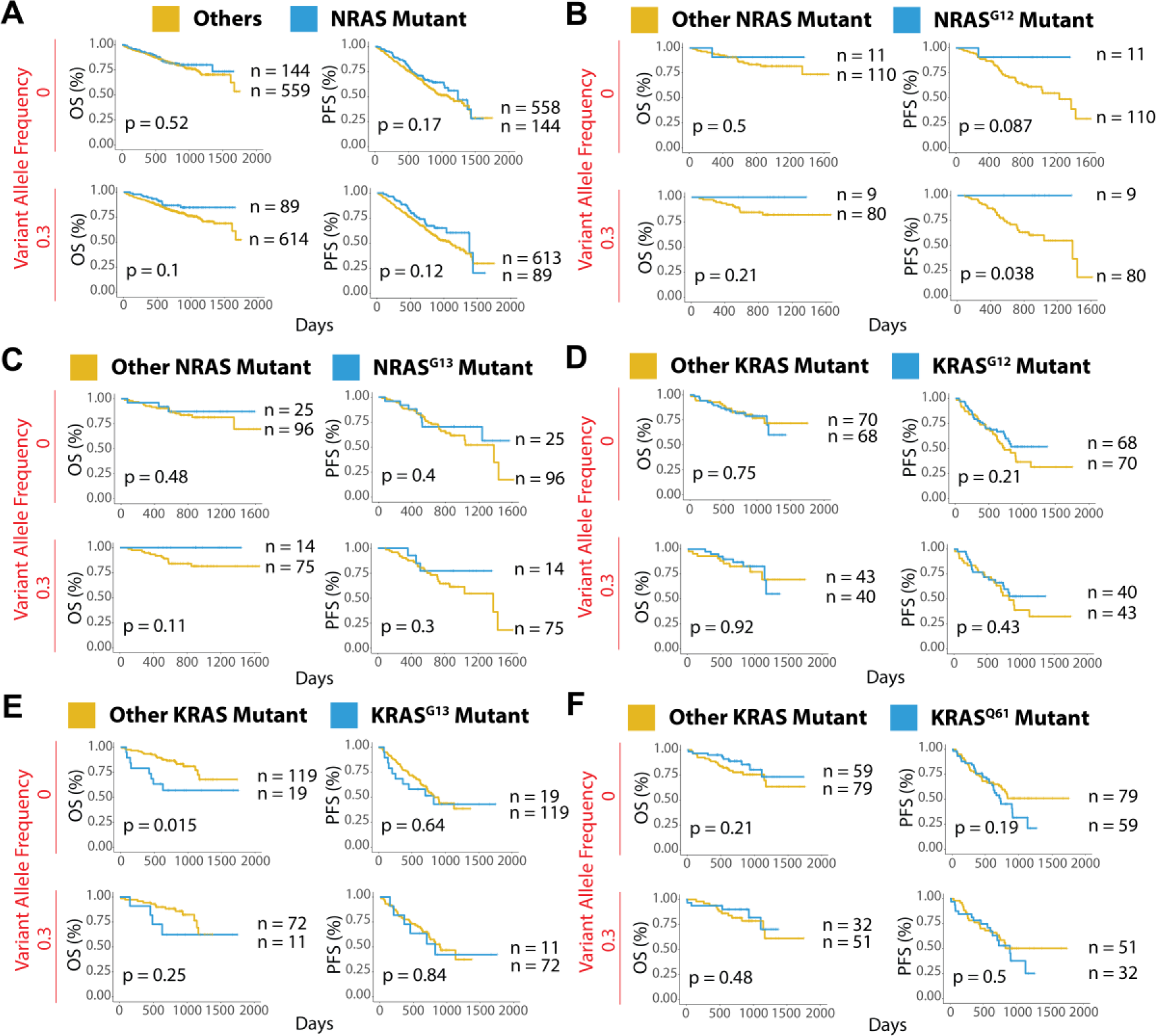
Survival analysis of MM patients with various *RAS* genotypes. **A.** Survival curves comparing the clinical outcome of newly diagnosed MM patients in the CoMMpass study with and without *NRAS* mutations. **B.** Survival curves comparing MM patients with codon-12 *NRAS* mutations versus other *NRAS* variants. **C.** Survival curves comparing MM patients with codon-13 *NRAS* mutations versus other *NRAS* variants. Finally, survival curves comparing MM patients with **D.** codon-12, **E.** codon-13, or **F.** codon-61*KRAS* mutations versus other *KRAS* variants. All analyses were performed at two levels of variant allele frequency cutoffs, 0 and 0.3, in order to assess the effect of clonality on survivorship. *p*-values are computed using log-rank tests.

**Figure S6.**
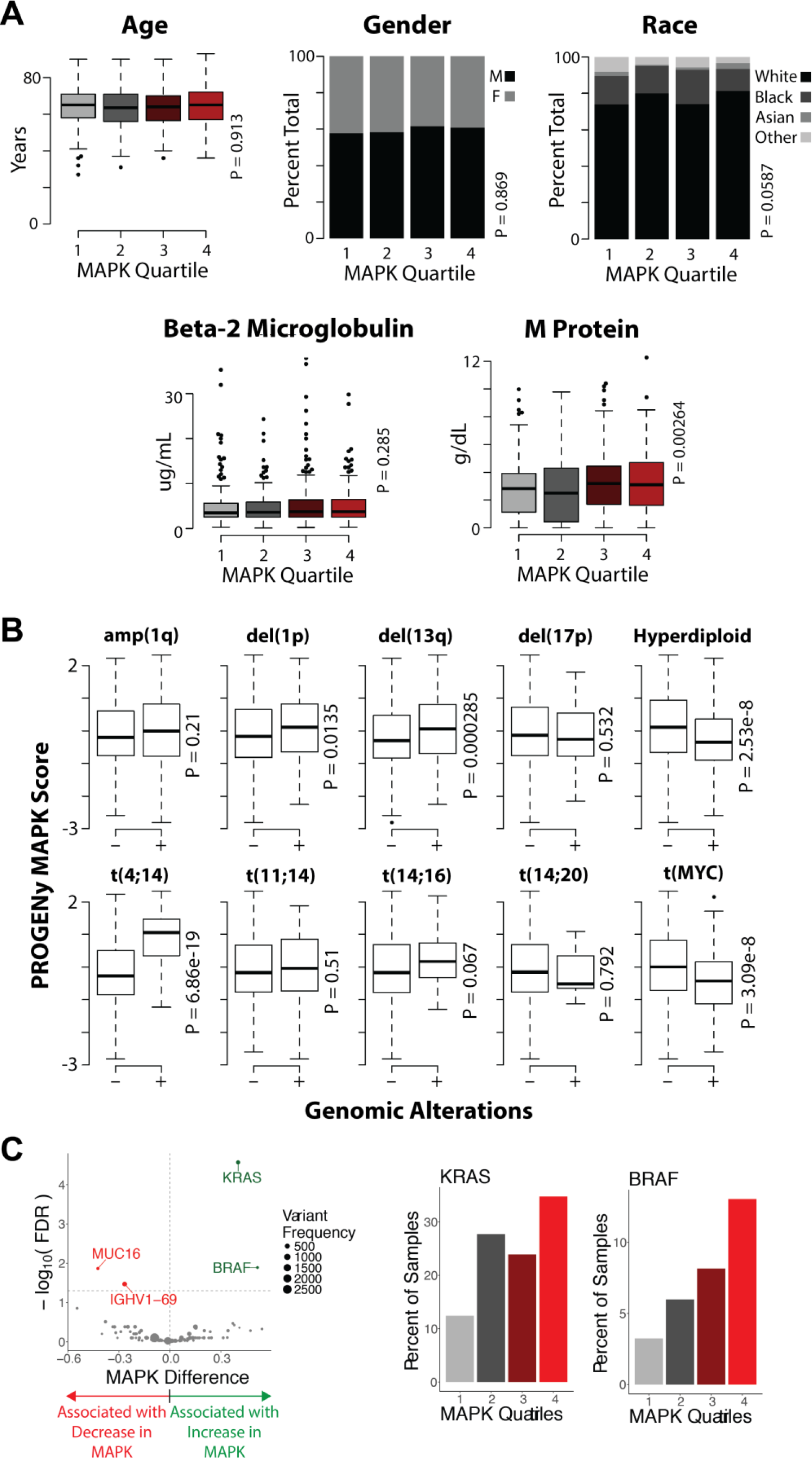
Association of PROGENy-predicted MAPK scores with demographic, clinical, and genomic features. **A.** Distribution of age, gender, race, beta-2 microglobulin levels, and M protein levels in MM patients are plotted against MAPK quartiles representing low (1) to high (4) predicted pathway activity. Only M protein levels are found to be linked to MAPK scores. **B.** Distributions of PROGENy-predicted MAPK activity for tumors with and without common genomic abnormalities identify signatures that are significantly associated with MAPK scores by Welch’s t-tests. These include del(1p), del(13q), hyperdiploidy, and t(MYC), with t(4;14) being the most strongly associated marker. **C.** Associations of the top 100 somatic mutations (from CoMMpass whole exome sequencing data) with predicted MAPK scores (from CoMMpass transcriptome data) in tumors from newly diagnosed MM patients show that mutations in *KRAS* and *BRAF* are significantly correlated with greater MAPK activity while tumors harboring variants of *MUC16* and *IGHV1-69* have significantly lower MAPK activity. The bar plots demonstrate that, in general, the fraction of patients with *KRAS* or *BRAF* mutations are greater in more activating MAPK quartiles. Welch’s t-tests on the average MAPK activity between mutated and non-mutated samples were used to determine significance. The *p*-values were adjusted for multiple comparisons by the Benjamini-Hochberg procedure.

**Figure S7.**
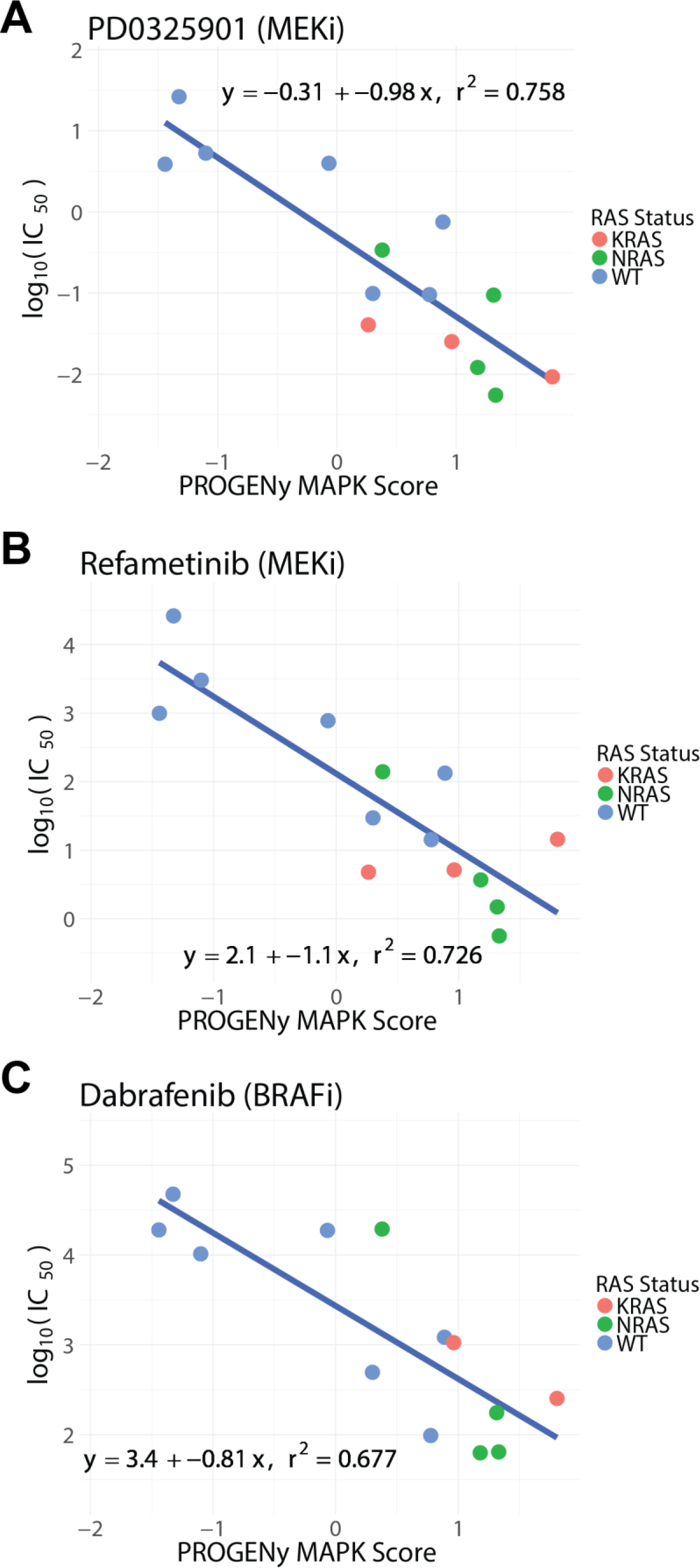
Correlation between sensitivity to small molecule inhibitors and PROGENy-predicted MAPK activation of myeloma cell lines. Sensitivities of MM cell lines to **A.** PD0325901 (MEKi), **B.** refametinib (MEKi), and **C.** dabrafenib (RAFi) show strong, linear relationships with predicted MAPK activity. Cell lines with greater predicted MAPK activation exhibit greater response to the three inhibitors, including some lines with WT *RAS* with greater sensitivity than *RAS* mutants. The line of best fit is displayed in blue along with its equation and coefficient of determination. Drug screen data from the Genomics of Drug Sensitivity in Cancer and RNAseq data from www.keatslab.org were used in the analysis.

## REFERENCES

1. Bolli, N., et al. Heterogeneity of genomic evolution and mutational profiles in multiple myeloma. Nat Comm 5, 2997 (2014).

2. Chapman, M.A., et al. Initial genome sequencing and analysis of multiple myeloma. Nature 471, 467–472 (2011).

3. Lagana, A., et al. Integrative network analysis identifies novel drivers of pathogenesis and progression in newly diagnosed multiple myeloma. Leukemia 32, 120–130 (2018).

4. Lohr, J.G., et al. Widespread genetic heterogeneity in multiple myeloma: implications for targeted therapy. Cancer Cell 25, 91–101 (2014).

5. Walker, B.A., et al. Identification of novel mutational drivers reveals oncogene dependencies in multiple myeloma. Blood 132, 587–597 (2018).

6. Casado, P., Hijazi, M., Britton, D. & Cutillas, P.R. Impact of phosphoproteomics in the translation of kinase-targeted therapies. Proteomics 17(2017).

7. Ruprecht, B. & Lemeer, S. Proteomic analysis of phosphorylation in cancer. Exp Rev Proteom 11, 259–267 (2014).

8. Casado, P., et al. Phosphoproteomics data classify hematological cancer cell lines according to tumor type and sensitivity to kinase inhibitors. Genome Biol 14, R37 (2013).

9. Casado, P., et al. Proteomic and genomic integration identifies kinase and differentiation determinants of kinase inhibitor sensitivity in leukemia cells. Leukemia 32, 1818–1822 (2018).

10. Dermit, M., Dokal, A. & Cutillas, P.R. Approaches to identify kinase dependencies in cancer signalling networks. FEBS Lett 591, 2577–2592 (2017).

11. McCormick, F. Targeting RAS directly. Ann Rev Biochem 2, 81–90 (2018).

12. Simanshu, D.K., Nissley, D.V. & McCormick, F. RAS Proteins and Their Regulators in Human Disease. Cell 170, 17–33 (2017).

13. Cox, A.D., Fesik, S.W., Kimmelman, A.C., Luo, J. & Der, C.J. Drugging the undruggable RAS: Mission possible? Nat Rev Drug Discov13, 828–851 (2014).

14. Li, S., Balmain, A. & Counter, C.M. A model for RAS mutation patterns in cancers: finding the sweet spot. Nat Rev Cancer 18, 767–777 (2018).

15. Newlaczyl, A.U., Hood, F.E., Coulson, J.M. & Prior, I.A. Decoding RAS isoform and codon-specific signalling. Biochem Soc Trans 42, 742–746 (2014).

16. Rasmussen, T., Kuehl, M., Lodahl, M., Johnsen, H.E. & Dahl, I.M. Possible roles for activating RAS mutations in the MGUS to MM transition and in the intramedullary to extramedullary transition in some plasma cell tumors. Blood 105, 317–323 (2005).

17. Liu, P., et al. Activating mutations of N- and K-ras in multiple myeloma show different clinical associations: analysis of the Eastern Cooperative Oncology Group Phase III Trial. Blood 88, 2699–2706 (1996).

18. Chng, W.J., et al. Clinical and biological significance of RAS mutations in multiple myeloma. Leukemia 22, 2280–2284 (2008).

19. Mulligan, G., et al. Mutation of NRAS but not KRAS significantly reduces myeloma sensitivity to single-agent bortezomib therapy. Blood 123, 632–639 (2014).

20. Billadeau, D., et al. Activating mutations in the N- and K-ras oncogenes differentially affect the growth properties of the IL-6-dependent myeloma cell line ANBL6. Cancer Res 57, 2268–2275 (1997).

21. Rowley, M. & Van Ness, B. Activation of N-ras and K-ras induced by interleukin-6 in a myeloma cell line: implications for disease progression and therapeutic response. Oncogene 21, 8769–8775 (2002).

22. Xu, J., et al. Molecular signaling in multiple myeloma: association of RAS/RAF mutations and MEK/ERK pathway activation. Oncogenesis 6, e337 (2017).

23. Soriano, G.P., et al. Proteasome inhibitor-adapted myeloma cells are largely independent from proteasome activity and show complex proteomic changes, in particular in redox and energy metabolism. Leukemia (2016).

24. Cox, J. & Mann, M. MaxQuant enables high peptide identification rates, individualized p.p.b.-range mass accuracies and proteome-wide protein quantification. Nat Biotechnol 26, 1367–1372 (2008).

25. R Development Core Team. R: A language and enviroment for statistical computing. R Foundation for Statistical Computing (2008).

26. Casado, P., et al. Kinase-substrate enrichment analysis provides insights into the heterogeneity of signaling pathway activation in leukemia cells. Sci Signaling 6, rs6 (2013).

27. Linding, R., et al. NetworKIN: a resource for exploring cellular phosphorylation networks. Nucleic Acids Res 36, D695–699 (2008).

28. Samatar, A.A. & Poulikakos, P.I. Targeting RAS-ERK signalling in cancer: promises and challenges. Nat Rev Drug Discov 13, 928–942 (2014).

29. Muller, E., et al. Pan-Raf co-operates with PI3K-dependent signalling and critically contributes to myeloma cell survival independently of mutated RAS. Leukemia 31, 922–933 (2017).

30. Way, G.P., et al. Machine Learning Detects Pan-cancer Ras Pathway Activation in The Cancer Genome Atlas. Cell Rep 23, 172–180 e173 (2018).

31. Guinney, J., et al. Modeling RAS phenotype in colorectal cancer uncovers novel molecular traits of RAS dependency and improves prediction of response to targeted agents in patients. Clin Cancer Res 20, 265–272 (2014).

32. Merchant, M., et al. Combined MEK and ERK inhibition overcomes therapy-mediated pathway reactivation in RAS mutant tumors. PloS ONE 12, e0185862 (2017).

33. Croonquist, P.A., Linden, M.A., Zhao, F. & Van Ness, B.G. Gene profiling of a myeloma cell line reveals similarities and unique signatures among IL-6 response, N-ras-activating mutations, and coculture with bone marrow stromal cells. Blood 102, 2581–2592 (2003).

34. Xu, J., et al. Dominant role of oncogene dosage and absence of tumor suppressor activity in Nras-driven hematopoietic transformation. Cancer Discov 3, 993–1001 (2013).

35. Tsherniak, A., et al. Defining a Cancer Dependency Map. Cell 170, 564–576 e516 (2017).

36. Steinbrunn, T., et al. Mutated RAS and constitutively activated Akt delineate distinct oncogenic pathways, which independently contribute to multiple myeloma cell survival. Blood 117, 1998–2004 (2011).

37. Walker, B.A., et al. Mutational Spectrum, Copy Number Changes, and Outcome: Results of a Sequencing Study of Patients With Newly Diagnosed Myeloma. J Clin Oncol 33, 3911–3920 (2015).

38. Schubert, M., et al. Perturbation-response genes reveal signaling footprints in cancer gene expression. Nat Comm 9, 20 (2018).

39. Kalff, A. & Spencer, A. The t(4;14) translocation and FGFR3 overexpression in multiple myeloma: prognostic implications and current clinical strategies. Blood Cancer J 2, e89 (2012).

40. Katoh, M. & Nakagama, H. FGF receptors: cancer biology and therapeutics. Med Res Rev 34, 280–300 (2014).

41. Yang, W., et al. Genomics of Drug Sensitivity in Cancer (GDSC): a resource for therapeutic biomarker discovery in cancer cells. Nucleic Acids Res 41, D955–961 (2013).

42. Ghobrial, I.M., et al. TAK-228 (formerly MLN0128), an investigational oral dual TORC1/2 inhibitor: A phase I dose escalation study in patients with relapsed or refractory multiple myeloma, non-Hodgkin lymphoma, or Waldenstrom’s macroglobulinemia. Am J Hematol 91, 400–405 (2016).

43. Rodrik-Outmezguine, V.S., et al. Overcoming mTOR resistance mutations with a new-generation mTOR inhibitor. Nature 534, 272–276 (2016).

44. Needham, E.J., Parker, B.L., Burykin, T., James, D.E. & Humphrey, S.J. Illuminating the dark phosphoproteome. Sci Signaling 12(2019).

45. Steinbrunn, T., et al. Combined targeting of MEK/MAPK and PI3K/Akt signalling in multiple myeloma. Br J Hematol 159, 430–440 (2012).

46. Wong, K.Y., et al. Frequent functional activation of RAS signalling not explained by RAS/RAF mutations in relapsed/refractory multiple myeloma. Sci Rep 8, 13522 (2018).

47. Burgess, M.R., et al. KRAS Allelic Imbalance Enhances Fitness and Modulates MAP Kinase Dependence in Cancer. Cell 168, 817–829 e815 (2017).

48. Bielski, C.M., et al. Widespread Selection for Oncogenic Mutant Allele Imbalance in Cancer. Cancer Cell 34, 852–862 e854 (2018).

49. Gupta, V.A., et al. Bone marrow microenvironment-derived signals induce Mcl-1 dependence in multiple myeloma. Blood 129, 1969–1979 (2017).

50. Burd, C.E., et al. Mutation-specific RAS oncogenicity explains NRAS codon 61 selection in melanoma. Cancer Discov 4, 1418–1429 (2014).

51. Hornbeck, P.V., et al. PhosphoSitePlus, 2014: mutations, PTMs and recalibrations. Nucleic Acids Res 43, D512–520 (2015).

52. McFarland, J.M., et al. Improved estimation of cancer dependencies from large-scale RNAi screens using model-based normalization and data integration. Nat Comm 9, 4610 (2018).

53. Meyers, R.M., et al. Computational correction of copy number effect improves specificity of CRISPR-Cas9 essentiality screens in cancer cells. Nat Genet 49, 1779–1784 (2017).

54. Cancer Cell Line Encyclopedia, C. & Genomics of Drug Sensitivity in Cancer, C. Pharmacogenomic agreement between two cancer cell line data sets. Nature 528, 84–87 (2015).

55. Love, M.I., Huber, W. & Anders, S. Moderated estimation of fold change and dispersion for RNA-seq data with DESeq2. Genome Biol 15, 550 (2014).

56. Gentleman, R.C., et al. Bioconductor: open software development for computational biology and bioinformatics. Genome Biol 5, R80 (2004).

57. Barwick, B.G., et al. Multiple myeloma immunoglobulin lambda transolcations portend poor prognosis. BioRxiv (2018). https://doi.org/10.1101/340877

58. Rausch, T., et al. DELLY: structural variant discovery by intergrated paired-end and split-read analysis. Bioinformatics 28, i333–i339 (2012).

